# SCORPy: Lowering the computational barrier to reproducible multiplexed imaging spatial single-cell proteomics analysis

**DOI:** 10.64898/2026.08.24.746722

**Authors:** Zoé Gerber, Samuel Simard, Harshitha Kolipaka, Zacharie Drouin, Juliane Sévigny, Violaine Pourcel, Celia del Carmen Crespo Oliva, Benjamin Tate, Korina Mouzakitis, Morgane Placet, Dominique Jean, Kiaya Deuel, Elias Pavlatos, Elizabeth Sturgill, Joanna Pucilowska, Gordon B Mills, Marilyne Labrie

**Author notes:** Corresponding author: Marilyne Labrie Dept Immunology and Cell Biology FMSS - Université de Sherbrooke 3201, Rue Jean-Mignault, Sherbrooke, QC, J1E 4K8.

## Abstract

Spatially resolved single-cell proteomic imaging technologies, including cyclic immunofluorescence (CycIF), generate high-dimensional data, critical for tissue-scale biological analysis. However, single-cell analysis remains computationally demanding, lacks standardization across platforms and is often inaccessible to experimental biologists without programming expertise. Here we present SCORPy (Single-Cell proteOmics Research Platform), a standalone, cross-platform desktop application that provides an end-to-end, code-free workflow for the analysis of single-cell proteomic data extracted from imaging experiments. SCORPy introduces methodological advances for preprocessing multiplexed imaging data: an exposure-aware, cycle-matched background correction strategy, and a normalization framework that harmonizes signal distributions across markers while enabling batch correction across experiments. These approaches are integrated with quality control, interactive thresholding and cell phenotyping using a hierarchical cell reference library, and downstream compositional and spatial analyses within a unified interface. Sample-level metadata can be incorporated throughout the workflow to support integrative analyses and facilitate generation of publication-ready visualizations. By combining robust preprocessing methods with an accessible implementation, SCORPy reduces computational barriers and promotes broader adoption of spatial single-cell proteomics analysis.

## Introduction

Recent advances in spatial proteomics and multiplexed imaging technologies^1^ have revolutionized this field, to the extent that spatial proteomics was named *Method of the Year 2024* by *Nature Methods*^2^. These technologies generate highly multiplexed, single-cell–resolved datasets while preserving tissue architecture and spatial context. Among them, multiplex cyclic-immunofluorescence (CycIF)^3^, COMET Lunaphore^4^ or CODEX PhenoCycler^5,6^ generate multiplex images and high-dimensional single-cell protein expression datasets. These approaches can be applied to both whole tissue sections and tissue microarrays (TMAs), producing datasets that quantify dozens of protein markers across hundreds of thousands of cells. These datasets are particularly informative for studying tumor cell state, tumor microenvironments, immune organization, neighborhood analysis, and tissue compartmentalization because they preserve both cellular phenotype and spatial position.

Although many single-cell spatial proteomic platforms have their own integrated registration and cell segmentation pipelines, analyzing the resulting single-cell data remains a major bottleneck, particularly for wet-lab researchers and clinicians who often lack computational expertise. While numerous analytical tools exist, most require proficiency in programming languages such as R or Python, limiting accessibility and slowing biological interpretation. Rigorous preprocessing, including background correction, normalization, and batch correction is essential prior to biological interpretation. Furthermore, cell phenotyping often requires defining marker-expression thresholds and constructing hierarchical classification rules, tasks that traditionally rely on custom scripting and programming expertise. Several tools address specific components of this workflow: CytoMAP^7^ emphasizes spatial analysis; SCIMAP^8^ provides a Python-based spatial single-cell analysis framework; CyLinter^9^ focuses on CycIF quality control; and pipelines like MCMICRO^10^, SPACEc^11^ or Tribus^12^ delivers end-to-end image-processing solutions, but do not rely on ground-truth expert knowledge. Unfortunately, these solutions either target isolated analytical steps or require command-line and programming proficiency. To our knowledge, no freely available platform provides a unified, end-to-end, code-free pipeline that bridges single-cell datasets to advanced spatial analysis within an integrated graphical environment. Such a platform would accelerate research by streamlining data analysis and lowering technical barriers for wet-lab researchers and clinicians who generate these valuable datasets but may lack computational expertise.

To address this gap, we developed SCORPy (Single-Cell proteOmics Research Platform), a code-free graphical user interface (GUI)-based desktop platform for the comprehensive processing and analysis of multiplexed imaging datasets. Unlike conventional image-processing pipelines, SCORPy focuses on the analysis of single-cell datasets generated from multiplexed proteomic imaging experiments. It accepts raw or pre-processed cell-level data from platforms such as CycIF, PhenoCycler, and COMET and provides a comprehensive framework for downstream phenotypic, cellular composition, and spatial analyses. SCORPy delivers a complete, guided analytical workflow encompassing data import, quality control, data filtering, background subtraction, data normalization, interactive cell phenotyping using a hierarchical cell reference library, marker-level analysis, cell quantification (proportions and densities), and spatial analysis (including grid-based with proximity mapping and neighborhood analysis). Importantly, the platform is compatible with data collected from whole tissue sections, tissue sections containing multiple regions of interest (ROIs), as well as TMAs where each tissue core is treated as a distinct ROI, enabling per-core analysis and across-core comparisons. SCORPy was designed around four principles: (1) vendor-agnostic data intake, (2) explicit and reproducible preprocessing logic, (3) portable and interpretable rule-based phenotyping, and (4) built-in spatial quantification that can be executed without programming expertise.

Here we present SCORPy and evaluate its methodological performance using a reference control TMA (ctTMA) dataset. The platform requires no coding and no software installation beyond running the executable and provides real-time visual feedback at every step.

## Results

### SCORPy end-to-end workflow overview

Multiplexed imaging technologies each present distinct advantages and limitations; here, we focus on CycIF, which enables visualisation and quantification of over 60 proteins through iterative cycles of staining and imaging. This acquisition workflow includes autofluorescence image acquisition, followed by sequential staining with primary antibodies labeled with fluorescence molecules, imaging and quenching cycles, multiplex image generation through image registration, and extraction of single-cell data through cell segmentation. The resulting single-cell data tables comprise for each cells the marker intensities, spatial coordinates, and metadata (Fig.1A). The resulting datasets can be processed and analysed using SCORPy (Fig1.B), a standalone cross-platform desktop application developed in Python that provides an intuitive, code-free graphical interface. Users can perform data preprocessing, cell phenotyping, and downstream analyses through interactive modules, while established computational methods are executed transparently in the background.

**Figure 1.**
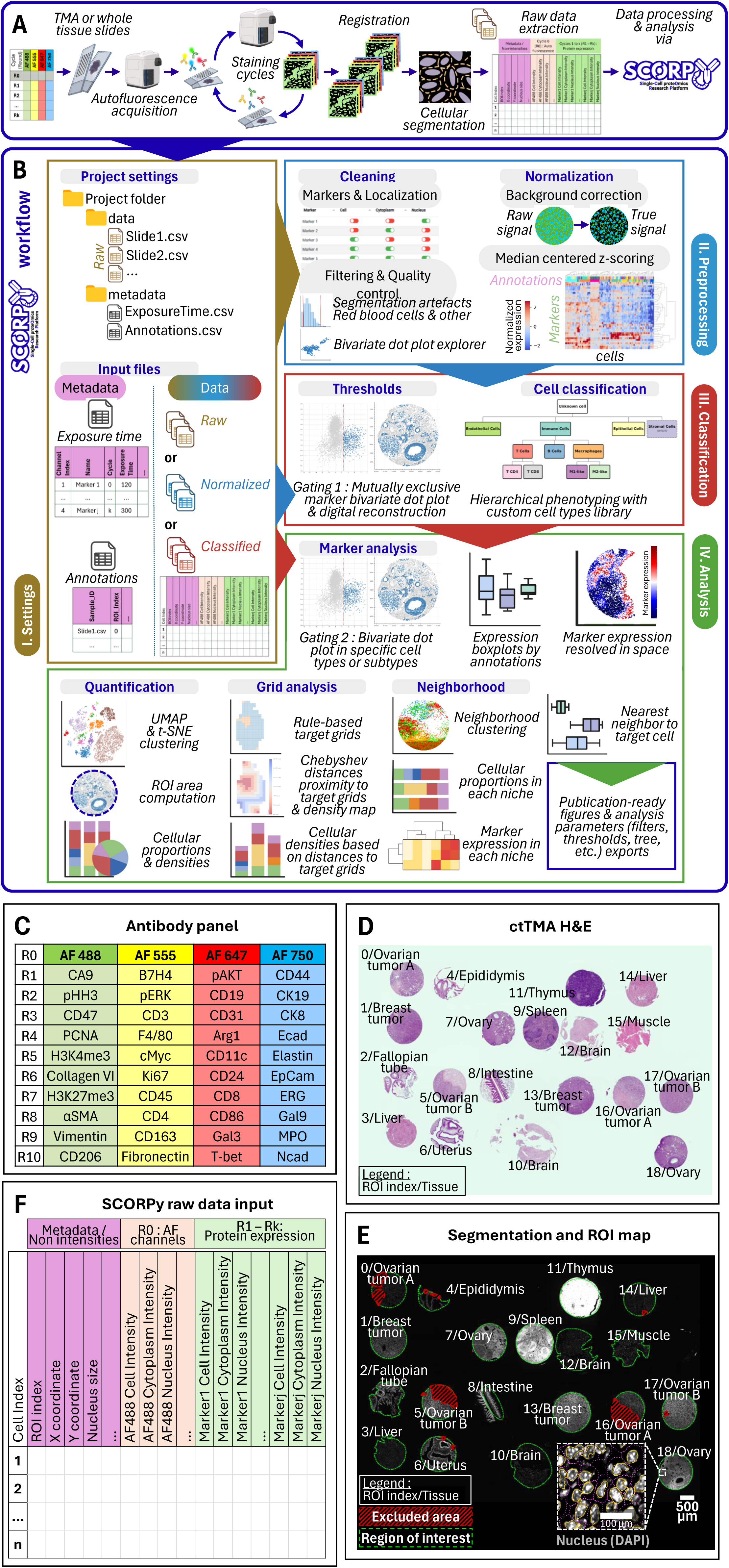
SCORPy end-to-end workflow. **(A) CycIF workflow overview.** Tissue sections from TMAs or whole-slide samples are subjected to iterative staining cycles using a designed antibody panel, including an initial empty round to assess autofluorescence in each channel, followed by sequential image acquisition. Images are then processed through registration, artefact identification and exclusion, cell segmentation, and extraction of single-cell measurements to generate raw tabular datasets containing protein marker intensities, spatial coordinates, morphometric features, and sample metadata. Single-cell datasets are subsequently imported into SCORPy for data harmonization, preprocessing, quality control, and downstream single-cell and spatial analyses. **(B) SCORPy analytical workflow.** Input data, including raw, normalized, or pre-classified single-cell tables and associated metadata, are organized within a standardized project structure. SCORPy accepts single-cell data tables exported from diverse image analysis segmentation platforms in comma- or semicolon-separated formats. The pipeline comprises quality control, data cleaning and localization assessment, and normalization (including background correction and median-centered z-scoring). Cells are classified using user-defined marker thresholds based on a mutually exclusive gating strategy, followed by hierarchical phenotyping. Downstream analyses include marker expression profiling, spatial mapping, cell-type quantification, grid-based spatial analysis, and neighborhood characterization. Publication-ready figures can be generated and exported along analysis parameters. **(C) CycIF antibody panel used to generate the reference ctTMA dataset.** Forty protein markers were distributed across 10 sequential staining cycles following an initial autofluorescence acquisition round. Markers were assigned to four fluorophore channels (AF488, AF555, AF647, and AF750) in each cycle, enabling iterative high-dimensional protein imaging and single-cell characterization of normal and malignant tissues. **(D) Representative H&E-stained section of the reference ctTMA.** The ctTMA contains normal and malignant murine tissues derived from multiple tissue types. Each tissue core was annotated as a region of interest (ROI). **(E) Representative ROI annotation and segmentation map of the ctTMA.** Tissue cores were manually delineated as independent regions of interest prior to image registration, cell segmentation, and downstream single-cell quantification. Segmented (green) and excluded (red) regions are shown, along with an inset illustrating a representative segmentation mask. Only nuclear staining (DAPI) is displayed in this immunofluorescence image. Scale bars: 500 µm. **(F) Representative raw single-cell data matrix generated following image registration and cell segmentation.** Each row represents an individual cell, with columns containing protein expression measurements (green columns), autofluorescence values (orange columns), and associated metadata (purple columns), including spatial coordinates and morphometric features. This standardized data structure serves as the primary input for downstream data analysis using SCORPy.

SCORPy structures the downstream analytical process into four interconnected modules: **Settings, Preprocessing, Classification, and Analysis**, that together form a guided and reproducible workflow (Fig.1B). The **Settings** module handles project path configuration, data import, and sample-associated metadata integration. The **Preprocessing** module performs quality control, outlier filtering, exposure time-aware background correction, and median-centered z-score normalization to standardize signal across markers and experiments. The **Classification** module enables interactive thresholding and a hierarchical decision tree-based cell library with visual feedback in both expression space and digital reconstructed tissue images. Finally, the **Analysis** module provides quantitative and spatial characterization, including unsupervised clustering analyses, cell population proportions, density estimation, grid-based spatial mapping, proximity analysis and neighborhood clustering. Together, these modules transform raw single-cell tables into reproducible, interpretable biological outputs (tables and publication-ready figures) within a unified and accessible framework tailored for experimental researchers. Importantly, SCORPy does not enforce a fixed analysis pipeline; users may enter the workflow at any stage by importing raw, pre- processed, or processed single-cell datasets and immediately access the relevant downstream analyses.

When initiating the pipeline, users can define their working directory, which must contain a *data/* subfolder with their raw CSV files and a *metadata/* subfolder with the exposure times of each marker, and optional annotation files. Output directories are then automatically created (*init/*, *cleaning/*, *normalization/*, *classification/*, *quantification/*, *spatial_analysis/* and *figures/*) for structured result storage. Multiple CSV files can then be selected and merged into a unified dataset. Before merging, the Settings module performs parallel pre-validation and quality control of all selected files using concurrent header inspection, checking for duplicate columns, empty files, and encoding errors, so that issues are caught before the full merge operation begins. An adaptive processing strategy automatically selects the optimal merging method based on file characteristics (number of files, total size, column compatibility), with real-time progress feedback during the operation. After merging, a comprehensive data quality report is generated, assessing for example NaN/null prevalence and column consistency across files.

Optional metadata import includes: (i) exposure times generated by registration tools such as ASHLAR^13^, or manually-generated, required for background subtraction; and (ii) annotation files containing clinical or experimental metadata (e.g., treatment group, patient ID, tissue type). Annotation files are loaded with automatic detection of join columns (Sample_ID and ROI_index), previewed in a paginated table, and merged with the dataset via a left join. Users select which annotation columns to include, and the merge result reports the number of matched and unmatched rows, providing immediate feedback on annotation coverage. An example of Exposure Time CSV file is available in Table S1.

### Data generation from a reference control TMA

To demonstrate the performance of SCORPy, we constructed a reference control TMA (ctTMA) composed of a diverse set of murine normal and cancer tissues. We evaluated SCORPy using the resulting multiplexed CycIF dataset comprising 40 protein markers measured across the 19 TMA tissue cores. Tissue microarrays enable parallel analysis of multiple samples under identical experimental conditions, thereby reducing staining variability and batch effects across experiments. While TMAs provide efficient and cost-effective multiplexed analysis, they inherently sample a limited tissue area, which may restrict the assessment of intratumoral heterogeneity. The extent of sampling depends on core size, which typically ranges from 0.6 to 2 mm in diameter (Fig.S1). To mitigate this limitation, ctTMA design incorporated multiple cores per tissue type and representative regions when available. The 40 antibodies were pre-conjugated to Alexa Fluor dyes (AF488, AF555, AF647, and AF750) or spectrally equivalent fluorophores, and distributed across 10 sequential staining rounds, following an initial autofluorescence acquisition (R0) (Fig.1C). Each ctTMA core was defined as a ROI (Fig.1D-E). Following image acquisition and registration, cell segmentation was conducted. Tissue folds and other technical artefacts, including regions affected by uneven or inconsistent staining, were excluded from pre-segmentation (Fig.1E). Segmentation generated a raw single-cell dataset comprising a total of 204,091 cells with per cell autofluorescence and marker intensities, spatial coordinates, and associated metadata (Fig.1F).

### Single-cell dataset importation and preprocessing

To support reliable downstream analysis, SCORPy Settings module includes an automated harmonization step when datasets are imported. During this step, the software validates file structure, detects and standardizes synonymous column names (for example sample and ROI identifiers), and parses multiple intensity-column conventions used across multiplex imaging technique exports into a common internal format. Metadata tables are also standardized by mapping common aliases, checking required fields, and filtering entries to those that match columns present in the imported data. Finally, annotation tables are merged using mandatory sample and ROI join keys to preserve row-level consistency. Together, these procedures allow heterogeneous single-cell datasets from different platforms and experimental designs to be integrated into a unified structure for consistent analysis. The complete list of recognized aliases for columns names is provided in Table S2.

Next, as multiplexed fluorescence imaging data are subject to several sources of technical variability, including autofluorescence, segmentation artefacts, and differences in exposure times across channels and imaging cycles, we included a multi-step Preprocessing module. First, given that most cellular segmentation tools can extract protein signals from specific compartments, users can restrict analysis to biologically relevant subcellular compartments using a marker localization selection step (Fig.2A). Then, quality-control filters are applied to remove residual segmentation artefacts, such as cells with extreme nuclear sizes, indicative of under-segmented cell aggregates. Additional filtering can be applied to exclude cells exhibiting abnormally high autofluorescence, commonly associated with red blood cell-rich regions, or other residual imaging artifacts that may persist despite prior tissue quality control (Fig.2B).

**Figure 2.**
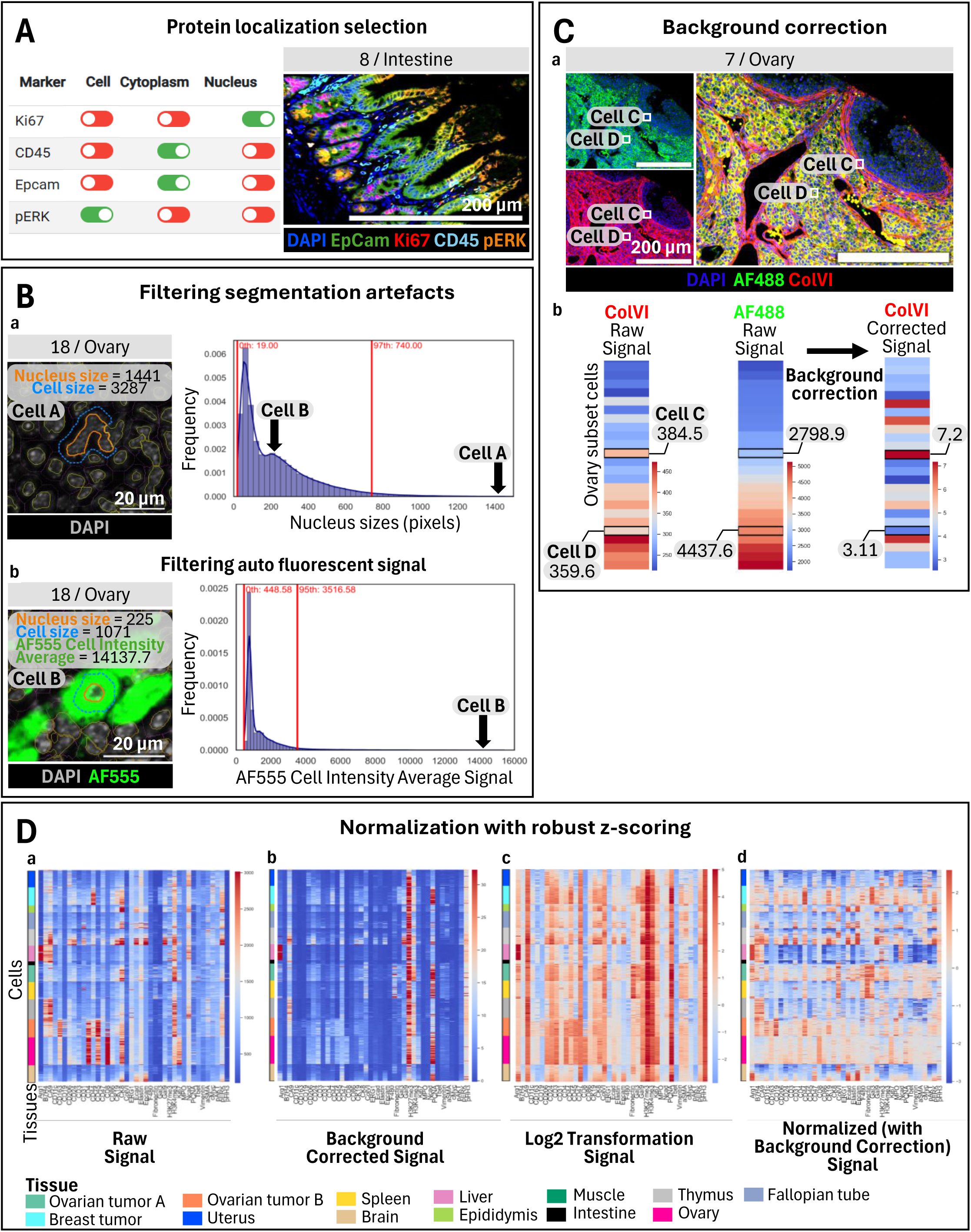
Preprocessing standardizes single-cell intensities and reduces technical variability. **(A) Marker localization selection.** Biologically relevant subcellular compartment can be selected for each marker prior to downstream analysis. Protein measurements extracted from the whole cell, nucleus, or cytoplasm/membrane can be interactively selected through a graphical interface, allowing compartment-specific quantification based on marker localization. **(B) Quality-control: Data filtering of segmentation and imaging artefacts.** Histogram of (a) the distribution of nuclear size and (b) autofluorescence intensities are used for quality-control assessment. Cells exhibiting extreme nuclear sizes, indicative of segmentation errors or under-segmented cell aggregates, and cells with abnormally high autofluorescence signals are excluded to minimize technical artefacts and improve dataset quality. As an example, Cell A represents two merged cells incorrectly segmented as one, while Cell B, despite accurate segmentation, is excluded due to elevated autofluorescence. Scale bars: 20 µm. Units: pixels. **(C) Background correction using autofluorescence signal.** (a) Ovary tissue images and (b) quantitative profiles illustrate subtraction of autofluorescence from Collagen VI (ColVI) protein signal using the corresponding autofluorescence channel (AF 488). This process reduces nonspecific fluorescence and improves discrimination between true biological signal and background. As an example, Cells C and D exhibit similar raw ColVI intensities; however, after correction, Cell C retains true signal while Cell D while Cell D is confirmed to lack ColVI expression. **(D) Normalization.** Heatmaps representing the marker expression across ctTMA tissue types at the single-cell level and at successive preprocessing stages, including (a) raw signal, (b) background correction, (c) log₂ transformation, and (d) post-normalization.

We next implemented an exposure time-aware background correction strategy that subtracts compartment-matched autofluorescence signals normalized by exposure time (Table S1). Specifically, marker and autofluorescence intensities are first normalized to their respective exposure times. The normalized autofluorescence signal from the corresponding channel is then subtracted from each marker intensity on a per-cell basis. This correction markedly improves signal distributions, particularly for tissues with high autofluorescence. For example, in the ctTMA dataset, Collagen VI (Col VI), detected in the AF488 channel, exhibited apparent positive signal in regions with strong autofluorescence (Fig. 2C). While these cells appeared Col VI-positive in the raw single-cell data, inspection of the corresponding images revealed that this signal was driven by high AF488 autofluorescence rather than true staining. By subtracting the exposure-normalized AF488 autofluorescence from the Col VI measurements, these non-specific signals were effectively removed. As a result, the corrected data no longer misclassified autofluorescence-high cells as Col VI-positive, and the resulting expression patterns showed improved biological concordance with the underlying tissue morphology.

The final step of data preprocessing involved log₂ transformation followed by median-centered z-score normalization of marker expression across all cells in the dataset (Fig. 2D). Specifically, for each marker, expression values are centred on the median and scaled by the standard deviation, reducing the influence of outliers while preserving relative differences between cells. Compared to log₂ transformation alone, this normalization reduced inter-marker variance and enabled direct comparison of expression levels across markers by placing them on a common scale^14^. As a result, high and low expression values become more interpretable across channels, facilitating threshold-based downstream analyses. Importantly, the inclusion of reference tissues spanning an expected range of marker expression can provide a reference for signal scaling, improving the distinction between high and low expression states across markers.

### Normalization using cross-experiment ctTMA reduces batch effects

Antibody-based, semi-quantitative techniques are inherently sensitive to batch effects arising from technical variability in staining efficiency, imaging conditions, and acquisition parameters across experiments. Consequently, shifts in signal intensity distributions are expected, even when protocols and reagents are nominally identical. To address this challenge, the Preprocessing module incorporates a normalization strategy based on the inclusion of shared reference tissues, such as ctTMA, within each experiment. By providing a common biological reference spanning a wide range of marker expression levels, the ctTMA enables calibration of signal intensities across datasets.

Log_2_ transformation alone does not resolve the scale incomparability between markers: across a panel of 20 to 60 proteins, each marker occupies a distinct dynamic range and population-level median, precluding direct cross-marker comparisons and rendering any single threshold value marker-specific rather than biologically generalizable. We therefore applied a modified z-score normalization, centering on the population median (*x̃*) and scaling by the population standard deviation, independently for each marker (column-wise), placing all markers on a common, dimensionless scale. The use of the median as the centering method is specifically designed to reduce the impact of extreme outliers that can occur when working with tissue samples. Different methods can be applied to normalizing and scaling single-cell proteomics data. We have found that log transformation combined with column-wise z-scoring achieve optimal results because it enables inter-cell comparisons and threshold transferability across ROIs and tissue types within a single experiment (Fig.2D, Fig.3C-E, Fig.S5C-E). To assess quality of the normalization, user-parameterized single-cell heatmaps with optional hierarchical clustering and multi-annotation sidebars allow visual comparison of raw and normalized data distributions, facilitating assessment of normalization quality before proceeding to cell phenotyping (Fig.2D, Fig.3C, Fig.S5C).

**Figure 3.**
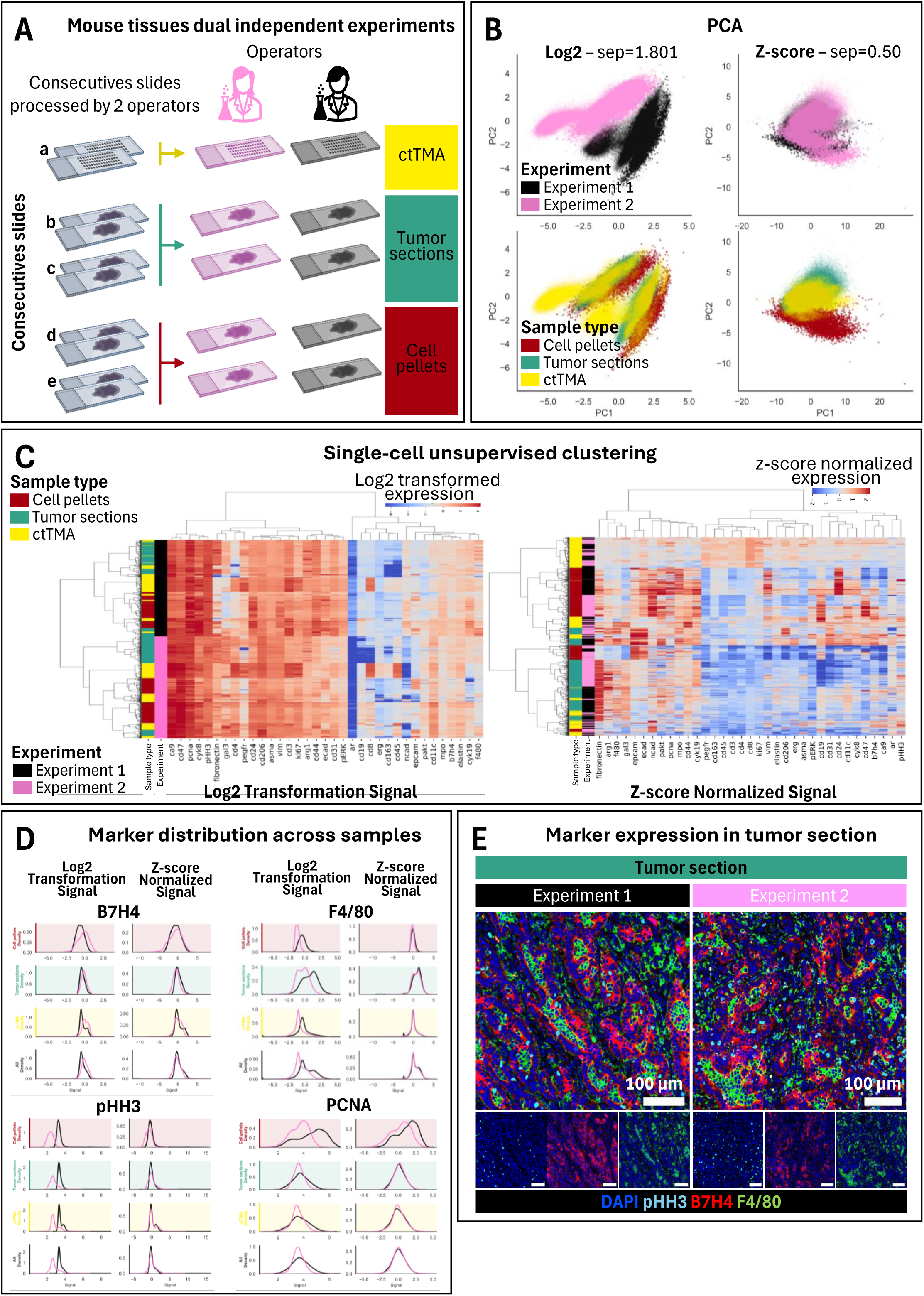
Cross-experiment normalization reduces batch effects while preserving biological structure. **(A) Experimental design for dual independents experiments.** Two consecutive sections of a (a) ctTMA, (b, c) two ovarian tumor tissue sections, and (d, e) two cell pellets were processed in two independent CycIF experiments conducted independently by two operators using an identical antibody panel. ctTMA sections served as biological reference samples to enable cross-experimental calibration and to assess technical reproducibility. **(B) Principal component analysis (PCA) before and after normalization by z-score.** PCA of single-cell data colored by experiment (top) and sample type (bottom). Log2-transformed data show strong separation between experiments (sep = 1.801), indicative of batch effects. After z-score normalization, separation between experiments is markedly reduced (sep = 0.50), while clustering by sample type (cell pellets, tumor sections, ctTMA) is preserved. **(C) Hierarchical unsupervised clustering heatmaps.** Heatmaps of marker expression across single cells following hierarchical unsupervised clustering, shown for log2-transformed (left) and z- score normalized (right) data. Cells are annotated by sample type and experiment. z-score normalization reduces experiment-driven clustering while maintaining biologically meaningful grouping of cell populations. **(D) Representative marker intensity distributions before and after ctTMA-based normalization.** Density distributions of selected markers (B7H4, F4/80, pHH3, and PCNA) across cell pellets, tumor sections, ctTMA reference, and the combined dataset while preserving overall expression patterns. Log2-transformed signals show variability between experiments, while normalization reduces batch effect. **(E) Representative marker expression in tumor tissue sections acquired in the two independent experiments.** Multiplexed IF images of tumor section from both experiments. Composite images show consistent spatial patterns of marker expression (DAPI, pHH3, B7H4, F4/80) across experiments. Insets highlight individual channels, demonstrating preservation of biological signal. Scale bars: 100 µm.

As an example, we analyzed two independents CycIF experiments performed on consecutive tumor sections and cell line pellet slides, processed by different operators using the same antibody panel (Fig.3A). Consecutive slides of both the experimental tumor samples and ctTMA were used to minimize variability in tissue composition between experiments. For batch correction, normalization parameters were derived exclusively from the ctTMA, using its median and standard deviation. This reference-based normalization enables consistent calibration across experiments while preventing biological differences in the experimental samples from confounding the normalization step. Principal component analysis (PCA) revealed strong batch separation in log2-transformed data with a separation coefficient of 1.8. In contrast, median-centered z-score normalization substantially improved mixing between experiments, reducing the separation coefficient to 0.5 (Fig.3B). Importantly, tissue-specific biological identities were preserved after normalization: hierarchical unsupervised clustering shows that samples continued to cluster according to tissue type rather than by experiment (Fig.3C). Marker distribution analysis further confirmed that normalization reduced technical variability while maintaining expression patterns across samples (Fig.3D–E, Fig.S6).

We further validated this approach using an independent hTMA dataset processed under similar conditions (Fig.S5A). As observed in the ctTMA analysis, normalization markedly reduced batch effects, with the PCA separation coefficient decreasing from 2.242 to 0.098 (Fig.S5B). Tissue-specific organization was maintained after normalization, with clustering driven by biological identity rather than experimental origin (Fig.S5C). Furthermore, marker distributions showed improved overlap following normalization compared to log₂ transformation alone (Fig.S5D–E). These results demonstrate that using control tissues for the normalization can effectively reduce batch effects without compromising biological interpretation of the data.

### Data processing: building a cell phenotype library

A major strength of single-cell analysis lies in the ability to assign cell identities and perform phenotype-resolved analyses based on co-expression of multiple markers. However, in multiplexed imaging datasets, defining marker positivity remains challenging, as robust and automated thresholding strategies are lacking, and signal distributions vary across markers and experiments. Semi-automated approaches, such percentile-based thresholds or clustering-assisted gating^15,16^ , are often used in practice to guide marker classification. While these methods can facilitate analysis, they remain sensitive to parameter choices, data quality, and user input, and frequently require manual adjustment. As a result, they can introduce variability and limit reproducibility across datasets and operators. Moreover, a key limitation of CycIF and related multiplexed imaging approaches is that they are typically applied to complex, multilayered tissues, where signal from individual cells can be contaminated by neighbouring cells due to limited spatial resolution and optical spillover. This can compromise accurate quantification of marker expression at the single-cell level and complicates the identification of discrete cell populations. To address this challenge, we implemented an interactive tool inspired by flow cytometry gating strategies and the RESTORE^17^ approach, enabling data-driven definition of marker thresholds. Marker expression cut-offs are defined using bivariate scatter plots, prioritizing mutually exclusive markers, when possible, to facilitate separation of positive and negative populations (Fig.4A). In parallel, a digital reconstruction of the tissue is generated, allowing users to iteratively refine thresholds by directly comparing the spatial distribution of marker-positive cells with the original fluorescence image of the tissue. This dual visualization approach ensures that gating decisions remain consistent with underlying tissue morphology.

**Figure 4.**
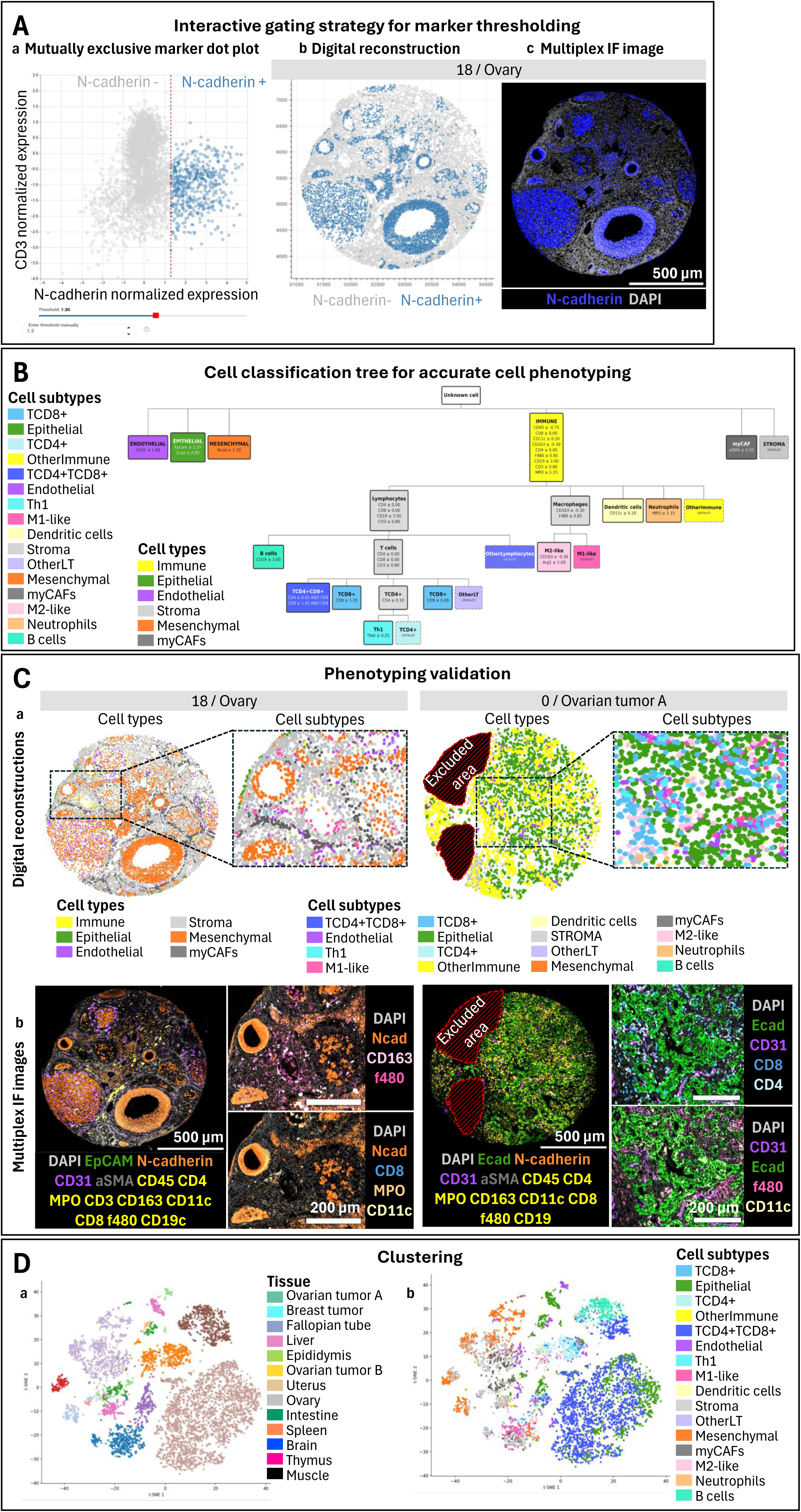
Hierarchical decision-tree phenotyping. **(A) Interactive marker gating and thresholds definition.** Flow cytometry-inspired gating strategy that enables data-driven and reproducible definition of marker expression thresholds. Thresholds are interactively adjusted using (a) bivariate scatter plots of normalized marker intensities. In parallel, a (b) synchronized digital tissue reconstruction provides real-time spatial visualization of marker-positive and marker-negative cells, allowing direct validation of gating decisions within their tissue context. This dual-view framework supports rapid, iterative refinement of thresholds while ensuring consistency between quantitative signal and tissue morphology. N-cadherin gating is shown as an example, alongside the (c) corresponding multiplex immunofluorescence image for visual reference. Scale bars: 500 µm. **(B) Hierarchical cell classification using a phenotyping decision tree.** A rule-based classification tree assigns cells to phenotypic populations through the sequential application of marker thresholds. Starting from all segmented cells, major compartments are first defined as broad cell types (e.g., immune, epithelial, stromal), followed by progressive subdivision into finer subtypes, including T cell subsets (CD4+, CD8+, Th1), B cells, macrophage states (M1-like, M2-like), dendritic cells, neutrophils, etc. In this hierarchical framework, terminal nodes (leaves) correspond to specific cell subtypes (e.g., epithelial cells, B cells, CD4+ T cells), whereas higher-level nodes define broader cell types. For analyses at the cell type level, only the first level of the hierarchy is considered; for example, all downstream immune subpopulations are grouped under the “IMMUNE” cell type. **(C) Validation of phenotypic annotations through digital tissue reconstruction.** (a) Digital tissue reconstructions illustrate cell classification results at both the cell type and cell subtype levels in representative samples (left: normal ovary; right: ovarian tumor A). Zoomed-in regions highlight the spatial organization and consistency of annotations across scales. Regions with insufficient data quality or artefacts were excluded from analysis. (b) Corresponding multiplex immunofluorescence images are shown to validate phenotypic assignments against marker expression patterns. This comparison demonstrates concordance between computational classification and tissue morphology, supporting the accuracy of the hierarchical phenotyping approach. This iterative validation process facilitates adjustment of gating thresholds and classification rules while ensuring concordance between computational annotations and the underlying biological signal. **(D) Clustering of single-cell phenotypes.** Two-dimensional embedding of single-cell data (t-SNE) colored by (a) tissue of origin and by (b) assigned cell subtype . Cells cluster according to both biological origin and phenotypic identity, indicating that the hierarchical classification preserves meaningful biological structure while enabling consistent annotation across diverse tissues.

A cell phenotype library is then created by the user through a hierarchical decision-tree framework that applies the gating strategies as explicit logical rules (Fig.4B). At the top level, major cellular compartments are defined by the user, using canonical lineage markers: for example, the ctTMA dataset was used to identify endothelial cells (CD31⁺), epithelial cells (EpCAM⁺ or E-cadherin⁺), mesenchymal cells (N-cadherin⁺), and immune cells (CD45⁺ and lineage-specific markers including CD3, CD8, CD4, CD19, CD11c, CD163, F4/80 and MPO). Cancer-associated myofibroblasts (myCAFs) are identified based on αSMA expression, whereas cells lacking these lineage markers are classified as stromal. At a second level, the immune compartment is further resolved into biologically relevant subtypes using combinatorial marker expression. For example, T cells can be subdivided into CD4⁺, CD8⁺ T cells, or double-positive populations, with additional annotation of functional states such as T-bet⁺ Th1 cells. Myeloid populations can be partitioned into macrophage subsets, including M1-like and M2-like phenotypes, as well as dendritic cells and neutrophils, based on marker-specific signatures (Fig.4B).

Importantly, gating decisions can be visualized through a digital reconstruction that incorporates all cell phenotypes listed in the tree (Fig.4C), ensuring that observed signal patterns accurately reflect the image of origin and the underlying biology. The Analysis module further enables an interactive and iterative construction of the phenotyping decision tree, allowing users to refine marker thresholds and classification rules at any stage of the analysis. This flexibility facilitates adaptation to dataset-specific characteristics while preserving analytical transparency. Once defined, decision trees can be saved, shared and reused across experiments, providing a consistent framework for cell type annotation and improving reproducibility across studies.

Once the cell phenotype library is established, a dimensionality reduction can be applied for visualisation purpose, using UMAP or t-SNE clustering on the classified dataset (Fig.4D). In our example, we observed that cells formed distinct clusters corresponding to the tissue of origin, reflecting the strong heterogeneity across tissues. When the same embedding was colored by cell subtype, these clusters resolved into biologically meaningful populations, with cells of similar phenotypes grouping together across tissues. Major cellular compartments remained well structured, and finer organization within immune populations revealed coherent subclusters corresponding to T cell subsets and myeloid lineages. This consistency between marker-defined phenotypes and low-dimensional structure supports the validity and robustness of the gating strategy.

### Data analysis: sample composition and marker expression

Once the cell phenotypes are assigned through the Classification module, the analysis module can be used for quantitative comparison of cellular composition and functional cell states across samples. The module enables simultaneous quantification of relative cell-type proportions, absolute cell densities and marker expression within a unified framework (Fig.5). Importantly, this framework is designed to support comparisons both within a single TMA and across multiple slides, while seamlessly integrating sample-level annotations. Indeed, users can define annotation groups, such as tissue types, experimental conditions, or regions of interest, across all samples or individual TMA cores, and directly generate quantitative summaries and visualizations based on these groupings.

**Figure 5.**
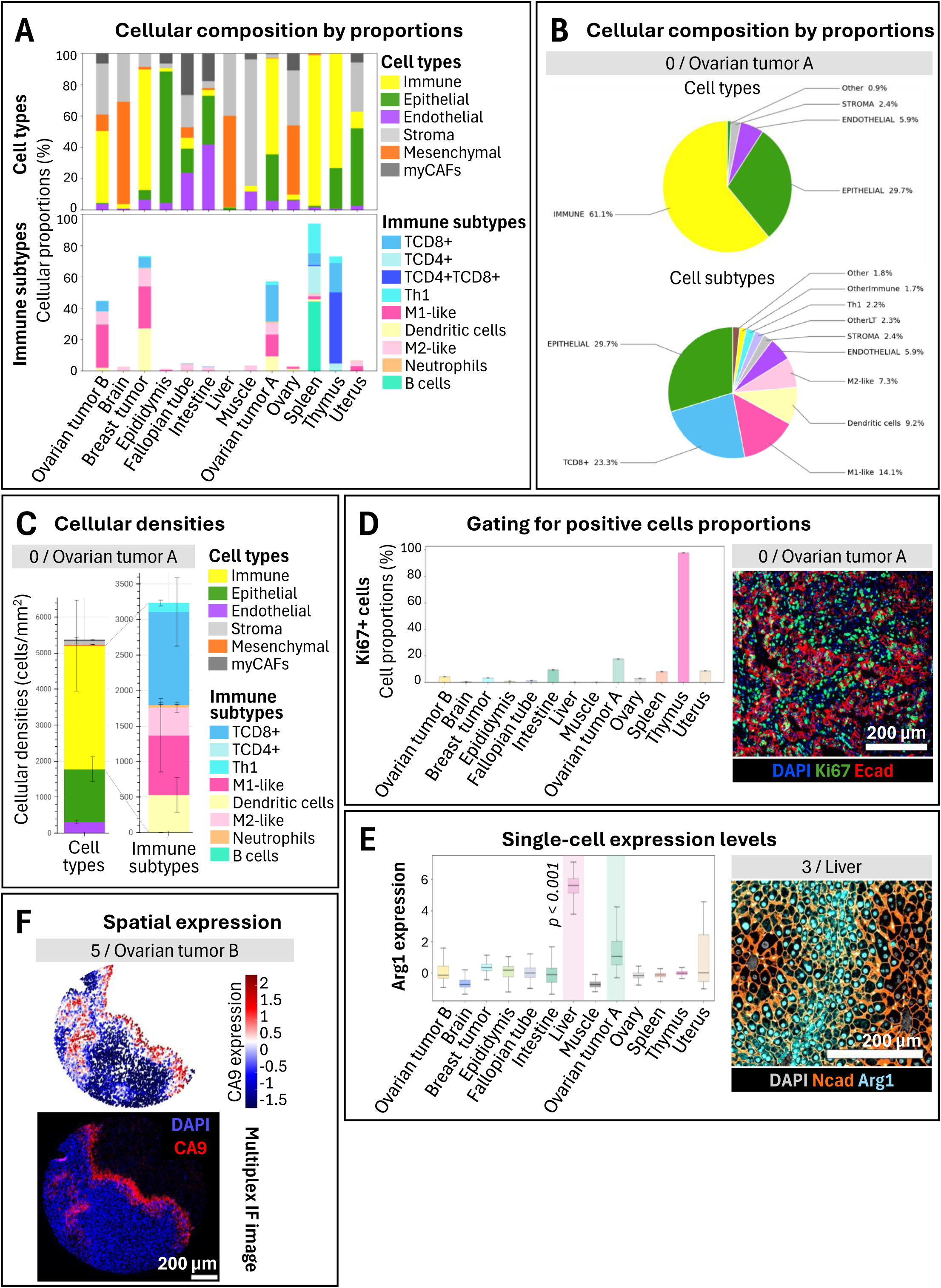
Quantitative characterization of sample-specific cell composition and marker expression. **(A) Cellular composition analysis across tissue types.** Quantitative comparison of cellular composition across samples, tissue types, experimental groups, or user-defined annotations. Representative stacked bar plots illustrate the relative abundance of major cell populations and subpopulations across ctTMA tissues, revealing tissue-specific cellular architectures and immune landscapes. **(B) Sample specific cell-type composition.** Pie charts summarize the percentage of cell types (top) and cell subtypes (bottom) in a representative sample (ovarian tumor A). This visualisation can be applied to a selected ROI, experimental groups, or any user-defined annotations and enables rapid characterization of tissue-specific cellular composition and immune infiltration patterns at the individual sample level. **(C) Cell density quantification.** Cell type (left) and immune subtype (right) densities (cells/mm²) measured in ovarian tumor A. **(D) Threshold-based marker expression analysis and spatial validation.** Proportion of Ki67- positive cells across tissues, reflecting differences in proliferative activity. A representative multiplex immunofluorescence image in ovarian tumor is shown alongside to illustrate marker expression and spatial distribution. Scale bars: 200 µm. **(E) Single-cell marker expression analysis.** Distribution of Arg1 expression across tissues at the single-cell level. Variations in expression highlight tissue-specific functional states, with representative imaging in liver tissue provided for validation. Scale bars: 200 µm. **(F) Spatial visualization of marker expression.** Continuous marker expression values are projected onto reconstructed tissue maps, allowing visualization of spatial expression gradients within the tissue context. Cells are colored according to marker intensity, revealing localized biological features and microenvironmental heterogeneity that may not be apparent from summary statistics alone. Spatial mapping of CA9 expression in a representative ovarian tumor sample, demonstrating hypoxia barrier. The corresponding multiplex image confirms localization patterns observed in the computational analysis. Scale bars: 200 µm.

As an example, using the ctTMA dataset, each core corresponds to a defined tissue type that can be analysed separately to capture tissue-specific differences in cell type composition. As shown in Fig.5A, epithelial tissues such as the uterus, epididymis or intestine show enriched epithelial cell populations, while lymphoid tissues such as the thymus or the spleen are enriched in immune cells. The immune cell subtypes analysis further shows that the thymus is primarily composed of CD4⁺CD8⁺ double-positive T cells, consistent with its role as a primary lymphoid organ supporting T cell maturation^18^, while the spleen is enriched in B cells, reflecting its role in humoral immunity^19^.

In tumor tissues, the immune compartment is much more heterogeneous, with high macrophage abundance spanning both M1-like and M2-like phenotypes. Importantly, the Analysis module allows the analysis of individual tissues. For example, Fig.5B shows the cell type and cell subtype composition of the ROI 0, corresponding to an ovarian tumor. This tumor displays extensive immune cell population, with more than 60% of cells classified as immune, including approximately 23% CD8⁺ T cells, consistent with the phenotype of an immune infiltrated tumor. These values can also be expressed as cell densities (Fig.5C), which is particularly useful when comparing cell abundance across samples. Importantly, the data associated to those figures can be exported as CSV files for downstream analysis at the end of the module.

The analysis module further enables flexible analysis of marker expression through complementary representations tailored to different biological questions. Marker expression can first be analysed using threshold-based binarization, where positivity is defined based on user-defined cut-offs. This allows direct comparison of the proportion of marker-positive cells across samples or conditions. For example, in the thymus, high Ki-67 positivity reflects active proliferation (Fig.5D). In parallel, the module also enables analysis of continuous expression values by aggregating single-cell measurements into distribution-based visualizations (box or bar plots) constructed from all cells within a selected sample or condition. Using this framework, we observed that Arg1 expression is significantly more elevated in liver tissue (ROI 3) compared to the other samples, highlighting tissue-specific metabolic activity (Fig.5E). Finally, we also integrated a spatial coordinates-based analysis to enable digital reconstruction of tissues, where continuous expression values are mapped onto cell positions using colour gradients. This representation allows visualization of spatially restricted expression patterns within the tissue context. For example, in an ovarian tumor, CA9 expression is localized to defined regions (Fig.5F), consistent with hypoxia-associated microenvironments.

### Data analysis: spatial analysis

A major strength of multiplexed imaging approaches is their ability to resolve cellular organization within intact tissue. However, most existing spatial analysis frameworks require computational expertise, limiting their accessibility. To facilitate spatial analysis, two complementary strategies have been implemented in the Analysis module: grid-based spatial analysis and neighborhood clustering (Fig.6).

**Figure 6.**
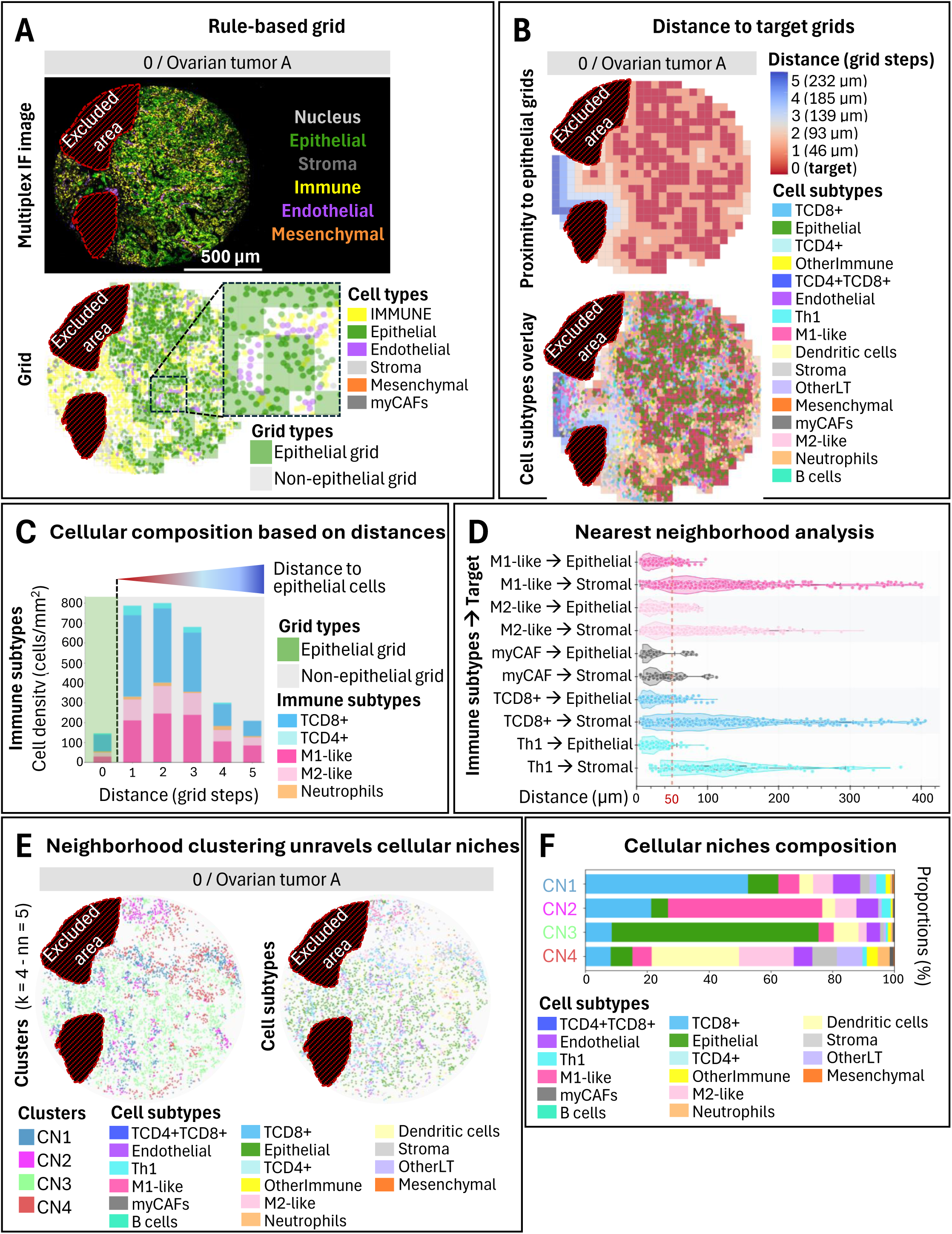
Spatial analysis reveals proximity-dependent organization of the tumor microenvironment. **(A) Grid-based spatial analysis framework.** Multiplex immunofluorescence image of a representative sample (ovarian tumor A) and its corresponding grid-based representation. Tissue sections are partitioned into regularly spaced grid units of user-defined size, generating a spatial map annotated according to local cellular composition. Grid units are classified based on the abundance of user-defined cell populations, enabling the identification of biologically relevant tissue compartments and spatial reference regions. In this example, grids are defined as epithelial or non-epithelial based on local epithelial cell content (threshold set at 50% within each grid). Excluded regions correspond to areas removed during preprocessing steps. **(B) Distance-to-epithelium mapping.** Following grid classification, distances are calculated relative to selected reference grids. Each grid unit is assigned to a discrete distance layer, enabling visualization of spatial relationships between tissue compartments and the generation of distance- resolved tissue maps. The heatmaps are showing the distance of each grid unit to the nearest epithelial region (expressed in grid steps and corresponding physical distances). Overlay of cell subtypes illustrates how different populations are distributed relative to epithelial structures, highlighting spatial gradients across the tissue. **(C) Distance-dependent cellular composition.** Quantification of cell-type abundance across successive distance layers from a reference compartment (epithelial regions), where layer 0 corresponds to the reference itself. This analysis characterizes spatial gradients in cellular composition and reveals how selected immune cell subtype densities vary as a function of distance to epithelial structures. **(D) Nearest-neighbor proximity analysis.** Distribution of distances between immune cell subtypes and their nearest epithelial or stromal neighbors. This analysis identifies preferential spatial associations, revealing subtype-specific interactions within the tumor microenvironment. A user-defined distance threshold in red (e.g., 50 µm) is applied to identify cells as being in close proximity and can be adjusted based on the analytical context. **(E) Neighborhood clustering identifies spatial cellular niches.** Unsupervised clustering of spatial neighborhoods defines distinct cellular niches (CN1–CN4). Spatial maps illustrate the distribution of these niches across the tissue alongside the contributing cell subtypes, revealing structured microenvironmental organization in ovarian tumor A. Clustering was performed with *k* = 4 clusters and *n*ₙₙ = 5 nearest neighbors. **(F) Composition of cellular niches**. Stacked bar plots showing the relative composition of each cellular niche in terms of cell subtypes. Each niche is characterized by a distinct combination of cell populations, reflecting functional heterogeneity within the tumor microenvironment. Notably, CN3 corresponds to an epithelial-enriched neighborhood, whereas CN4 represents a mixed immune cell niche.

In the grid-based approach, tissue sections are partitioned into regularly spaced regions of user-defined size, generating a spatial grid overlaid onto the tissue (Fig.6A). Each grid unit is then annotated based on its cellular composition. For example, grids can be classified as epithelial if they contain a defined proportion of epithelial cells (e.g., ≥50%), and as non-epithelial otherwise. This classification enables the definition of spatial reference regions within the tissue. Distances between grid units are subsequently computed relative to regions of interest, allowing each grid to be assigned to discrete distance bins. In this framework, epithelial grids are defined as distance 0, and surrounding grids are grouped into successive distance layers (Fig.6B). These distance-resolved maps can be visualized directly on digital tissue reconstructions, enabling intuitive interpretation of spatial relationships between cell populations. Using this approach, SCORPy identifies spatial gradients in immune cell infiltration. For example, CD8⁺ T cell density decreases as a function of distance from epithelial regions in an ovarian tumor, revealing a progressive decrease in density with increasing distance from epithelial grids (Fig. 6C). This analysis quantitatively captures tissue organization by linking cell-type abundance to proximity from biologically defined compartments. Importantly, this framework is flexible and can be applied to any cell type or feature of interest such as immune cell enriched, enabling systematic investigation of spatial interactions across tissues and conditions.

To complement the grid analysis, the Analysis module can also be used to perform nearest-neighborhood analysis between user-defined source and target cell populations (Fig.6D). Users define source (e.g. M1-like macrophage) and target cell populations (e.g. epithelial cells), and a violin plot is created where each point represents an individual source cell plotted according to its distance (µm) to the nearest target cell. As an example, Fig.6D shows distinct spatial preferences across immune populations. For example, in an ovarian tumor infiltrated by immune cells, CD8⁺ T, Th1, M1-like and M2-like cells are preferentially located near epithelial cell populations compared to stromal cells, consistent with immune surveillance at the tumor interface, whereas myCAFs showed no apparent proximity bias to either target’s populations. For visualization purposes, in the example, an arbitrary threshold of 50 µm was applied to define cells considered “close” to target populations and can be adjusted regarding the biological question.

To further characterize higher-order spatial organization within the microenvironment, we applied neighborhood clustering based on local cell-type composition (Fig.6E). This approach groups spatially proximal cells into discrete niches by leveraging the composition of their immediate microenvironment, rather than pairwise distances alone. Unsupervised clustering on the ovarian tumor A identified four well-defined cellular neighborhoods (k = 4 clusters, N = 5 neighbors) with distinct compositional signatures, highlighting the presence of spatially structured microenvironments within this tissue. Further quantification of niche composition (Fig.6F) demonstrates that these clusters are enriched for specific combinations of immune, stromal, and epithelial populations. For instance, some niches were characterized by a predominance of epithelial and CD8⁺ T cells (CN1), consistent with immune-enriched tumor interfaces, whereas others were dominated by mixed immune compositions, including CD4⁺ T cells, dendritic cells, and B cells (CN4), suggesting functionally diverse immune microenvironments. Together, these spatial analysis functions allow analysis of tissue architecture and organization into distinct cellular niches with characteristic compositions, reflecting spatially coordinated cellular interactions.

### Performance across multiplex spatial proteomics platforms and analytical workflows

To assess the compatibility of SCORPy across multiple multiplexed imaging platforms, we analyzed independent datasets generated using COMET and PhenoCycler technologies. First, a TMA was independently stained, imaged, and segmented using the Lunaphore COMET platform by the Immune Monitoring and Cancer Omics (IMCO) platform at Oregon Health & Science University. The resulting single-cell dataset was analyzed independently using SCORPy and the IMCO in-house analytical pipeline, providing a stringent benchmark of SCORPy against an established academic platform using distinct normalization, quality-control, and cell-classification workflows.

The raw dataset was imported into SCORPy through the Settings module and normalized using the Preprocessing module. The gating strategy and corresponding thresholds were established from mutually exclusive scatter plots (Fig.S3A). These parameters were subsequently used to assign each cell to a cell type according to a defined hierarchical classification tree in the Classification module. The corresponding cell-type assignments were visualized spatially on representative TMA cores (Fig.S3B). Despite being generated through independent computational approaches, both pipelines produced highly concordant spatial maps and cell-type distributions across representative TMA cores (Fig. S3B-F), demonstrating that SCORPy reproduces the biological conclusions generated by an independent expert analytical platform despite substantial methodological differences in filtering, preprocessing and cell classification. Importantly, methodological differences between the pipelines extended beyond normalization approaches to include distinct filtering and quality control procedures. Following removal of the AF555 signal, SCORPy retained around 35,000 cells from the original 43,000-cell dataset, whereas the IMCO pipeline maintained approximately 40,000 cells. This differential filtering likely reflects distinct quality control criteria and signal processing parameters between the two systems.

Quantitative analysis revealed a slightly higher proportion of classified immune cells in the SCORPy output compared to IMCO (48.9% vs. 46.7%). Overall, cell population frequencies were highly concordant between pipelines, with differences generally remaining below 5% for each cell populations. This discrepancy may be attributed to the high cellular density and potential cell overlapping in tissue samples, where marker signal spillover could affect neighboring cells, making thresholding more difficult. The minor variations in cell type proportions between pipelines can also be explained by operator-dependent manual gating of markers, which introduces variability in threshold determination (Fig.S3A). Minor differences were observed in the classification of specific immune subsets. The IMCO pipeline exhibited a higher percentage of non-classified immune cells (Fig.S3C), whereas SCORPy identified a greater proportion of M2-like macrophages with Δ=5.3% (Fig.S3D-F). Importantly, spatial inspection of the underlying multiplex images supported the presence of M2-like macrophages in several regions identified by SCORPy (Fig.S3E-F) but not by the IMCO pipeline, suggesting that SCORPy may provide improved resolution of specific cellular phenotypes in complex tissue environments.

To determine whether this performance extended beyond COMET-derived data, we next applied SCORPy to a publicly available PhenoCycler dataset generated using a distinct imaging technology and analytical workflow.(Fig.S4)^20^. This dataset comprises single-cell protein measurements for 57 antibody markers, along with corresponding cell segmentation and spatial coordinates. SCORPy enabled comprehensive analysis of tissue architecture, cellular neighborhoods, and regional variation along the intestinal axis. To assess regional variation in cell composition, we first computed cellular densities of stromal, immune, and epithelial subtypes across anatomical regions of the intestine, including small bowel and colon segments (Fig.S4A). Distinct spatial distributions of major cell compartments were observed, reflecting known tissue organization. Epithelial cells were enriched in mucosal regions, while immune cell abundance varied along the intestinal tract. Statistical analysis comparing the small bowel and colon revealed significant differences in the densities of various cell subtypes, underscoring the regional specialization of intestinal tissues. We utilized the quantification tab of the Analysis module to perform t-SNE clustering, which revealed clear segregation of major epithelial, immune, and stromal populations (Fig.S4B). Using the protein expression resolved in space section, we analyzed the localization of Ki67, a marker of proliferation, in the proximal jejunum of patient B011 (Fig.S4C). Ki67 expression was localized at the basal level of the proximal jejunum and was not expressed in immune cells. Multiplex CODEX images with Cytokeratins (green), Ki67 (red), and CD45 (cyan) further confirmed epithelial structures, proliferative zones, and immune cell localization, respectively. The spatial restriction of proliferative (Ki67+) cells to specific regions, such as crypt-like domains, supports the biological fidelity of the data and provides insights into the proliferative activity across tissue regions. To investigate higher-order spatial organization, we performed a cellular neighborhood analysis with parameters k=7 and nearest neighbors nn=3 (Fig. S4D). This analysis identified 7 distinct cellular neighborhoods (CN1– CN7) corresponding to recurring multicellular assemblies, such as epithelial-rich regions, immune-enriched niches, or mixed compartments. Comparison with underlying cell subtype maps showed that these neighborhoods capture higher-order tissue organization beyond individual cell identities, reflecting coordinated spatial patterning. These visualizations support the ability of the analytical framework to extract biologically meaningful spatial domains from CODEX/PhenoCycler data. Collectively, analyses of Cyc-IF, COMET, and PhenoCycler datasets demonstrate that SCORPy provides a technology-agnostic framework for multiplex spatial proteomics analysis, enabling consistent cell classification, spatial interrogation, and tissue architecture mapping across diverse imaging platforms.

## Discussion

SCORPy addresses a practical and methodological gap in multiplexed imaging analysis by providing a unified, end-to-end framework for single-cell spatial proteomics that remains accessible to non-computational users. While a growing ecosystem of tools exists for image-based analysis, these tools typically operate at different stages of the workflow and often require advanced technical expertise or fragmented pipelines to achieve full analysis.

Upstream image analysis platforms such as QuPath^21^ or CellProfiler^22^ can provide powerful solutions for image segmentation, feature extraction, and, in some cases, deep-learning-based cell classification. However, these tools generally focus on the image processing stage and require downstream integration with custom scripts or external platforms for phenotyping, quantification, and spatial analysis. In contrast, SCORPy operates downstream of segmentation and integrates data preprocessing, phenotype assignment, quantitative analysis, and spatial investigation within a single executable environment, thereby eliminating the need for multi-tool orchestration and reducing analytical complexity. Similarly, spatial analysis frameworks such as CytoMAP^7^ or SCIMAP^8^ offer advanced methods to interrogate spatial cellular organization, including neighborhood analysis, clustering, and spatial statistics. While powerful, these approaches typically require programming expertise and do not provide integrated workflows for data preprocessing or biologically guided cell phenotyping. SCORPy complements these tools by providing a fully integrated and flexible analysis pipeline, where spatial analysis is directly linked to user-defined phenotypes and sample metadata within an interactive interface. Recent efforts such as SPACEc^11^ or STELLAR^15^ further aim to streamline multiplexed image analysis through interactive Python-based workflows. While these approaches improve usability compared to fully scripted pipelines, they still require installation through command-line environments (e.g., Docker containers or Windows Subsystem for Linux), which may remain a barrier for non-computational users. In contrast, SCORPy is distributed as a standalone executable that can be directly deployed on local machines without programming or environment configuration, further lowering the barrier to entry for experimental researchers and helping bridge the gap between advanced spatial proteomics analytics and experimental researchers.

A key strength of SCORPy is its user-centered design, which enables researchers to perform complex analyses through an intuitive interface without requiring specialized computational expertise or dedicated software infrastructure. Despite its user-friendly design, SCORPy is capable of handling large-scale datasets, including millions of cells, hundreds of TMA cores, and whole-tissue sections, making it suitable for both exploratory studies and high-throughput translational applications. Importantly, the platform supports flexible data import and export at all stages of the pipeline, allowing users to incorporate sample-level metadata, reorganize input formats with minimal effort (e.g., column header adjustments), and generate high-resolution, publication-ready figures directly from the interface.

Rather than relying on fully automated classification, SCORPy adopts a human-in-the-loop analytical paradigm, enabling expert-driven phenotyping through interactive gating strategies inspired by flow cytometry. This design contrasts with black-box machine learning approaches and emphasizes interpretability, allowing users to iteratively refine thresholds based on both quantitative distributions and spatial validation within tissue reconstructions. While this approach enhances transparency and biological interpretability, particularly in heterogeneous tissue contexts where signal contamination, autofluorescence, and biological complexity can challenge fully automated methods, it does require user expertise and time investment. Incorporating optional semi-automated or guided strategies in future iterations could further balance interpretability with efficiency, enabling users to leverage automation where appropriate without compromising control.

Reproducibility and transparency are central to the SCORPy framework. All intermediate outputs can be systematically saved, creating a complete audit trail of the analysis. Phenotyping strategies are encoded as explicit decision trees that can be exported in both human-readable (PNG) and machine-readable (JSON) formats, facilitating reproducibility and reuse across experiments. Threshold values and analysis parameters can be exported and re-imported as CSV files, ensuring consistency across studies and enabling collaborative exchange of analytical workflows. While comprehensive output retention enhances transparency, it may introduce storage and memory considerations for very large datasets. To address this, future implementations may further emphasize lightweight reproducibility strategies, such as prioritizing the retention of parameterized workflows and decision logic over full intermediate datasets, thereby reducing storage overhead while preserving analytical traceability.

Importantly, SCORPy is designed to be platform-agnostic, supporting single-cell data derived from CycIF, Lunaphore, CODEX, and other multiplexed imaging technologies. While minor formatting adjustments (e.g., header standardization) may be required depending on the upstream pipeline, this process is straightforward and does not require programming expertise. This interoperability enables seamless integration into existing workflows and broad applicability across experimental platforms.

The normalization framework presented here demonstrates robustness to inter-experimental variability and supports cross-sample comparability across independent datasets, including consecutive ctTMA sections and distinct tissue types. Importantly, downstream compositional and spatial analyses remain biologically structured following preprocessing, supporting the validity of the approach in preserving meaningful biological signals while reducing technical variation.

Nonetheless, several limitations should be considered. First, SCORPy operates on pre-extracted single-cell data tables and does not perform image registration or segmentation, requiring users to rely on external tools for upstream processing and to refer back to the original images for validation. Second, the absence of cell-level ground truth annotations limits direct benchmarking of phenotyping accuracy, necessitating validation through consistency with immunofluorescence images and known biological patterns. Third, some spatial analyses implemented, such as grid-based approaches, simplify the underlying continuous tissue architecture, although this is mitigated by the inclusion of complementary methods such as nearest-neighbor analysis and spatial clustering. Finally, integration of data-driven phenotype discovery approaches remains important directions for future development, particularly in the context of rapidly advancing artificial intelligence and machine learning techniques.

In summary, SCORPy bridges the gap between complex computational workflows and biological interpretation while preserving methodological transparency or interpretability. By combining accessibility, flexibility, and reproducibility within a single framework, SCORPy enables broader adoption of multiplexed imaging analysis and supports consistent, scalable investigation of spatial cellular organization across diverse complex tissues.

## Methods

### Study design and datasets

#### ctTMA construction

All tissue specimens used in this study were collected and handled in accordance with institutional ethical guidelines and approved protocols. For the ctTMA and hTMA construction, formalin-fixed paraffin-embedded (FFPE) tissue samples were assembled into a single paraffin block using a tissue arrayer. Cylindrical cores (0.6 and 1 mm diameter) were extracted from donor FFPE blocks and arrayed into a recipient block (Fig.S1).

#### CycIF experiment

CycIF was performed as previously described^23,24^. Briefly, tissue sections were deparaffinized, and antigen retrieval was performed in pH 6 citrate buffer using a pressure cooker (Cuisinart CPC-600) for 20 minutes. Slides were rinsed in distilled water and further incubated in pH 9 Tris/EDTA buffer at elevated temperature for 15 minutes. Non-specific binding was blocked using phosphate-buffered saline (PBS) supplemented with 10% normal goat serum and 1% bovine serum albumin (BSA). Tissue autofluorescence was acquired prior to antibody staining using an Axioscan 7 fluorescence slide scanner (Zeiss). Sequential staining, imaging, and fluorophore inactivation cycles were then performed for a total of 10 cycles. Each cycle involved incubation with up to four primary antibodies conjugated to Alexa Fluor 488, 555, 647, or 750 fluorophores, followed by whole-slide imaging using the Axioscan 7 system. Fluorescence signals were subsequently quenched using a solution containing 3% hydrogen peroxide and 20 mM NaOH in PBS for 30 minutes. Complete signal inactivation was verified prior to subsequent staining cycles. A detailed list of antibodies is provided in Table S3. For unconjugated antibodies, labeling was performed using Alexa Fluor conjugation kits (Invitrogen: A37570, A37571, A37573, A37575) according to the manufacturer’s instructions.

#### Image processing and single-cell feature extraction

Raw CycIF images were processed using established computational tools. Image registration across staining cycles was performed using ASHLAR^13^. Cell segmentation, visualization, and feature extraction were carried out using QI Tissue Image Analysis software^25^ (version 1.4.0, Quantitative Imaging Systems). Mean fluorescence intensities for each marker were extracted within predefined cellular compartments (e.g., cell, nucleus, cytoplasm), depending on marker localization. The resulting single-cell dataset was then exported and used as input for downstream processing and analysis using SCORPy.

#### Lunaphore COMET dataset

A human TMA comprising multiple tissue types; triple-negative breast cancer, clear cell renal cell carcinoma, prostate cancer, normal lymph node, and endometrial cancer; was generated, sequentially stained, and segmented by the Lunaphore COMET™ 1.0 platform, as previously described^26^. The 19-plex panel included CCR2, CD103, CD11b, CD11c, CD163, CD20, CD3, CD4, CD45, CD68, CD8, FOXP3, granzyme B (GZMB), HLA class II, Ki-67, pan-cytokeratin (PanCK), PD-1, PD-L1, and α-smooth muscle actin (αSMA).

Multiplex stitched and aligned image was exported as OME-TIFF file and analyzed using Visiopharm Phenoplex™ software (version 2025.08.1.18881, 64-bit). Following image and tissue-quality assessment, nine TMA cores with adequate tissue integrity and staining quality were retained for analysis. Regions of interest were defined as each core, and a pretrained machine-learning algorithm was applied to the first-cycle DAPI image to identify analyzable tissue. Nuclei were segmented using Visiopharm’s built-in DAPI U-Net algorithm, followed by uniform expansion of the nuclear boundaries to approximate whole-cell regions. Fluorescence intensity was then quantified for each marker on a per-cell basis.

#### Lunaphore COMET analysis

Marker-positivity thresholds were established by manual gating for each core (ROI) independently to account for tissue-specific differences in staining intensity and background fluorescence. Gating results were exported to R (version 4.4.3), where a proprietary classification script was used to assign cellular phenotypes and quantified cell populations within each tissue compartment. Cell densities were calculated as the number of classified cells per unit area (cells/mm²). All downstream data processing, statistical analyses, and visualization were performed in R.

#### SCORPy analysis

The raw-intensity dataset was parallelly and independently processed and normalized with SCORPy. First, a quality-control filter was applied to remove cells exhibiting abnormally high autofluorescence. Then, data were normalized using SCORPy Preprocessing module.

The gating strategy and corresponding thresholds were then established from mutually exclusive scatter plots, as previously described. These parameters were subsequently used to assign each cell to a cell type according to a defined hierarchical classification tree.

#### PhenoCycler dataset

We performed a simple analysis of a publicly available PhenoCycler dataset of human intestinal tissues^20^ (Fig.S4). The dataset along with all additional information can be downloaded at https://datadryad.org/dataset/ doi:10.5061/dryad.pk0p2ngrf. As the data was already normalized, segmented and annotated, it was directly loaded into the Analysis module of SCORPy.

#### Software implementation and deployment

SCORPy was implemented in Python using the Panel framework^27,28^, with Bokeh^29,30^ for interactive visualization. The application follows a layered architecture: each analytical module is divided into a UI layer (widget layout and user interaction) and a Logic layer (data transformations and algorithms). Cross-module communication is achieved through a typed observer pattern, where modules subscribe to event classes (e.g., data availability changes, threshold updates) and receive notifications without direct coupling. Shared state is managed through thread-safe singletons. The platform operates as a local web application encapsulated within a native desktop window using pywebview^31,32^. This approach provides a seamless desktop experience without requiring users to interact with a web browser. It is distributed as a standalone executable built with PyInstaller 6.19.0^33,34^, requiring no Python installation or dependency management by the end user.

SCORPy v1.0.0 was developed with Python 3.13.12, Panel 1.7, pandas 2.3, NumPy 2.2, Bokeh 3.7, SciPy 1.15, Seaborn 0.13, Altair 5.5, scikit-learn 1.6, and UMAP 0.5. During development, the authors used AI-based Copilot coding assistant for code generation, refactoring, and debugging; all generated code was manually reviewed and tested before inclusion.

As researchers increasingly work with large-scale datasets, memory efficiency becomes a central consideration in the platform design. The platform employs a shared data store that transparently offloads intermediate DataFrames to disk as Apache Parquet^35,36^ files when not in active use, then reloads them on demand. This lazy-loading strategy reduces peak memory consumption during multi-step analyses. A real-time memory monitoring widget, displayed in the application header, continuously tracks RAM usage and alerts users when predefined thresholds (75%, 85%, and 92%) are reached. Input/output operations leverage the Polars^37^ library for parallelized multi-threaded CSV reading, immediate type optimization, and memory-efficient file merging via a Polars / Parquet / pandas bridge that avoids holding both representations in RAM simultaneously.

#### User interface organization

The user interface is organized into four modules that reflect the logical progression of the workflow:

**(I) Settings**, **(II) Preprocessing**, **(III) Classification**, and **(IV) Analysis** (Fig.1B). These modules are divided in eleven sequential tabs: (I.1) Instructions, (I.2) Project settings, (I.3) Input files, (II.1) Cleaning, (II.2) Normalization, (III.1) Thresholding, (III.2) Cell classification, (IV.1) Marker analysis, (IV.2) Quantification, (IV.3) Grid analysis and (IV.4) Neighborhood analysis. This modular structure guides users from data import and configuration through preprocessing, cell type assignment, and downstream quantitative and spatial interrogation, while maintaining clear separation between analytical stages.

The pipeline manages data through five processing states:

(1) *RAW* → *MERGED* → *CLEANED* → (2) *NORMALIZED* → (3) *CLASSIFIED*

Users may enter the pipeline at three stages depending on the state of their data: (1) raw post-segmentation data for the full pipeline, (2) pre-normalized data to skip directly to thresholds and classification (Classification module), or (3) pre-classified data to skip directly to marker analysis, quantification, or spatial analysis (Analysis module).

#### Input harmonization and schema standardization

SCORPy accepts CSV files (comma or semicolon-separated, auto-detected) containing per-cell measurements exported from cell segmentation tools. A typical input file includes, for each cell: an identifier for the sample, file, slide or image, an optional region of interest index (useful for TMA samples where each core corresponds to a ROI), spatial coordinates of the cell centroid or nucleus, nucleus or cell sizes, and fluorescence intensity measurements for each marker at each subcellular localization (Fig.1). Column name standardization is performed automatically through a regex-based format detection system that recognizes multiple naming conventions. The platform parses intensity column names using ordered regex patterns covering formats such as "Marker Localization Metric" (CycIF) or "Marker: Localization: Metric" (PhenoCycler), and underscore-delimited variants. Non-intensity annotation columns (e.g., Sample_ID, ROI_index, Nucleus_Size) are identified via an extensive alias list supporting over 15 common naming variants per column type (Table S2).

#### Quality control and filtering

Default filtering thresholds are set at the 95th percentiles (q∼0.95) for nucleus size and autofluorescence intensity, with adjustable paired or independent sliders. Additional custom filters can be applied to any numeric column. A bivariate dot plot explorer, inspired by the RESTORE method^17^, enables visualization of marker co-expression patterns to evaluate data quality before filtering. Automated cleaning removes DAPI columns, empty or unassigned intensity channels (e.g., r6c5), and columns not matching the selected markers and localizations.

#### Background signal correction and normalization

This step corrects raw fluorescence intensities by accounting for exposure time and subtracting cycle-matched autofluorescence. Each marker column is first divided by its own exposure time (retrieved from the metadata, if provided), and the corresponding autofluorescence column, matched by fluorophore position within the imaging cycle, is independently divided by its own exposure time before subtraction:

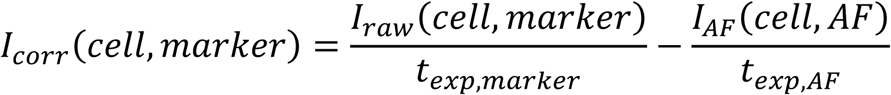

Where *I_raw_* is the raw fluorescence intensity, *t_exp_*_,*marker*_ and *t_exp_*_,*AF*_ are the respective exposure times for the marker and its matched autofluorescence channel, and *I_AF_* is the autofluorescence intensity corresponding to the same fluorophore position within each imaging cycle.

Background-corrected intensities are first log2-transformed (values ≤ 0 set to a floor of 0.1) to stabilize variance and symmetrize the right-skewed intensity distributions typical of fluorescence data before using median-centered z-scoring:

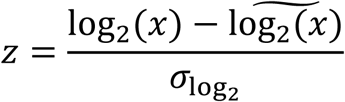

Where *σ* denotes the population standard deviation. When datasets exceed 100 MB, computation is parallelized across CPU cores using Dask^38^ partitioning.

#### Thresholding and marker positivity

The Thresholds tab enables users to define marker positivity thresholds through interactive Altair^39^ scatter plots, also inspired on the RESTORE mutually exclusive marker method^17^. For each marker, cells are displayed along the expression axis with an adjustable threshold slider. Cells with expression values above threshold are highlighted, and a paired Bokeh-based digital reconstruction plot shows the spatial distribution of positive cells within each sample and ROI, using the original cell or nucleus centroid coordinates. Saved thresholds can be exported and re-imported for reproducibility. Threshold definitions can be converted into binary positivity columns for downstream analyses.

The Marker Analysis tab extends the threshold visualization to classified data, enabling users to explore marker expression patterns filtered by cell type or cell subtype. By filtering cells according to their classification, users can assess how marker expression varies across biologically defined populations. Digital reconstruction plots can be colored by classification labels to reveal spatial distributions of specific populations. SCORPy also allows users to generate binary marker-positivity columns based on user-defined thresholds and cell-type constraints. These binary variables can then be used as inputs for downstream spatial grid analysis rules, enabling the study of spatial interactions between defined functional cell states. Alternatively, these derived variables can be exported for external statistical or bioinformatics analyses, facilitating integration with complementary analytical workflows.

#### Hierarchical cell phenotyping

The classification algorithm is fully vectorized using NumPy : (1) all marker threshold evaluations are pre-computed as boolean arrays in a single pass across the entire dataset; (2) the tree is traversed depth-first, left-to-right (priority order), where at each node marker conditions combined with AND/OR logic are evaluated against the pre-computed arrays; (3) the first matching node assigns the cell type (root-level children) or cell subtype (deeper levels); (4) cells matching no sibling node are assigned to a designated personalizable default node (e.g., "Stroma" or "Other").

#### Tissue area calculation

Two methods are supported: (a) using an existing area column from the dataset with unit conversion (mm², µm², px²) when available, or (b) estimating the occupied tissue area from cell centroid coordinates. In the latter method, a 2D histogram is computed over the cell positions using a user-configurable *B* number of bins per axis, dividing the spatial extent of detections into a *B* × *B* grid. The tissue area is then estimated as:

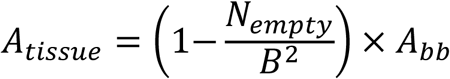

Where *A_bb_* = (*x_max_* − *x_min_*) × (*y_max_* − *y_min_*) is the bounding box area in pixels² computed from the extreme cell coordinates, *B*^2^ is the total number of histogram bins, and *N_empty_* is the number of bins containing zero cells. The ratio *N_empty_*/*B*^2^ estimates the fraction of the bounding box that is unoccupied (e.g., empty regions between TMA cores, tissue tears or background), and the complement gives the fraction covered by tissue. The resulting area in pixels² is converted to physical units using the imaging resolution *r* (in µm/pixel):

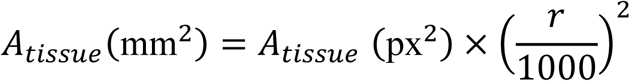

Increasing *B* provides a finer-grained estimate that excludes small empty regions, while lower values yield a coarser approximation.

#### Dimensionality reduction

Users select the embedding method (UMAP or t-SNE), choose which markers to include (all intensity markers by default or a user-selected subset), and configure algorithm-specific parameters. UMAP parameters include the number of neighbors (*k*, default: 15), minimum distance (default: 0.1), and distance metric (Euclidean, Manhattan, Chebyshev, or cosine). t-SNE parameters include perplexity (default: 30) and maximum iterations (default: 500). Both methods share a configurable random state for reproducibility, a cell sampling size (default: 5,000) to maintain interactive performance on large datasets, and optional filtering by any annotation column to restrict the embedding to specific subpopulations (e.g., only immune cells, or only a particular treatment group). When the number of marker features exceeds 15, a PCA pre-reduction step automatically reduces dimensionality to 30 components (or the available minimum) using randomized SVD before applying the embedding algorithm, improving both speed and separation quality.

#### Spatial grid and proximity analysis

For grid-based approach, the tissues are partitioned into rectangular grid cells using either: (a) a target-cells-per-grid-cell mode, where the grid dimensions adapt to maintain approximately *N* cells per grid cell, or (b) fixed pixel dimensions specified by the user. Each ROI is processed independently, making this approach naturally suited for TMA datasets where each core is analyzed in isolation. Grid cells are classified using an ordered, priority-based rule system. 4 types of rules can be defined: proportion rules (classify a grid cell if the proportion of cells matching a condition exceeds a user-defined threshold), binary column rules (classify based on the mean of a binary positivity column created in the Marker Analysis tab), expression rules (classify based on mean marker intensity) and Composite rules (combine multiple conditions with AND/OR logic). Rules are evaluated top-to-bottom with first-match semantics; grid cells matching no rule receive a configurable default label.

Proximity analysis computes the Chebyshev distance (8-connectivity) from every grid cell to the nearest source grid type a C-implemented distance transform. Physical distances are converted from grid steps to micrometers and millimeters based on imaging resolution. Proximity values can be injected as new columns into the classified dataset for downstream statistical analysis.

#### Spatial neighborhood analysis

Cell-resolution neighborhood analyses operate directly on segmented cell centroids. All neighborhood computations are performed per ROI independently and then aggregated at the result level, so that spatial proximity is never computed across discontinuous tissue cores. Cartesian nucleus coordinates (in pixels) are converted to physical units (µm) using a user-specified imaging resolution set to match the acquisition pixel size (default 0.346 µm per pixel), and all spatial queries use a k-d tree (‘scipy.spatial.cKDTreè) for efficient fixed-radius and nearest-neighbor lookups. Where a search radius is required, a single user-adjustable value (default 25 µm) is applied uniformly across all ROIs.

For each source and target cell label, defined by a selected cell type or cell subtype, the Euclidean distance to the nearest target cell within the same ROI is recorded. The query is unconditional (no distance cutoff): the user-specified radius is retained only as a visual reference line on the resulting plots, so that the full distribution of source-to-target distances is preserved. Distances are pooled across ROIs and displayed as per-pair horizontal violin distributions overlaid with the median, interquartile range, and Tukey whiskers, enabling comparison of how closely a given phenotype is positioned relative to candidate interaction partners.

To identify recurrent multicellular spatial motifs, each cell is assigned to a cellular neighborhood following the windowing approach of Goltsev et al^6^. For every cell, a composition vector is built by counting the cell-type identities of its ***N*** nearest neighbors (default ***N*** = **10**); using a fixed neighbor count rather than a fixed radius makes the vector robust to local variation in cell density. Composition vectors are **log**(**1** + ***x***)-transformed and pooled across all selected ROIs, and a k-means model (default ***k*** = **5**; random state = 42) partitions cells into CN clusters. Each CN is summarized by its mean cell-type composition profile (back-transformed and normalized to proportions), and cells are re-mapped to their tissue coordinates for spatial visualization of neighborhood organization.

#### Validation strategy in the absence of cell-level ground truth

Given the absence of pathologist-labeled cell-level ground-truth annotations, validation in this study relies on orthogonal consistency checks: marker-expression coherence within assigned classes, expected spatial organization patterns, and concordance with reference IF marker images in selected regions. Reference IF images used for this concordance assessment are available alongside the associated single-cell raw table and the parameter and tree JSON files used to generate classification and analysis outputs.

#### Statistical analysis

Group comparisons use non-parametric tests selected automatically by group number and pairing. For two groups, the Mann–Whitney U test (unpaired) or the Wilcoxon signed-rank test (paired) is applied; for more than two groups, a Kruskal-Wallis’s omnibus test (unpaired) or Friedman omnibus test (paired) is followed by pairwise Mann-Whitney or Wilcoxon tests. Pairwise p-values are adjusted for multiple comparisons using the Holm–Bonferroni procedure.

Effect sizes are reported as the rank-biserial correlation r. Groups with fewer than two observations are excluded from testing. Comparisons can be computed at the single-cell level or on per-sample aggregates (median per sample) to respect biological replication.

#### Exports and result storage

All analysis outputs, including the enriched dataset with cell type, cell subtype, binary marker columns, grid type, and proximity columns, can be exported as individual per-sample CSV files or a single merged CSV file. All figures will be saved by request by the user in the dedicated *figures/* directory.

#### Data availability

SCORPy software is freely available under the MIT license. Pre-built executables for Windows and macOS, wiki documentation, and ctTMA dataset alongside the corresponding fluorescence image in OME-TIFF format, are available at https://github.com/marylab26/SCORPy and https://doi.org/10.5061/dryad.d2547d8jn, enabling full reproducibility of the analyses.

## Acknowledgments

We thank the HORBITUS platform at the Université de Sherbrooke (https://www.usherbrooke.ca/medecine/recherche/notre-caractere-distinctif/infrastructure-et-plateformes-de-la-recherche/horbitus) for their support with the ctTMA sample preparation and CycIF experiments.

We also thank the Lunaphore COMET platform and the IMCO team at OHSU for their support with the human TMA sample preparation and COMET experiments.

---

**Supplementary Figure 1.**
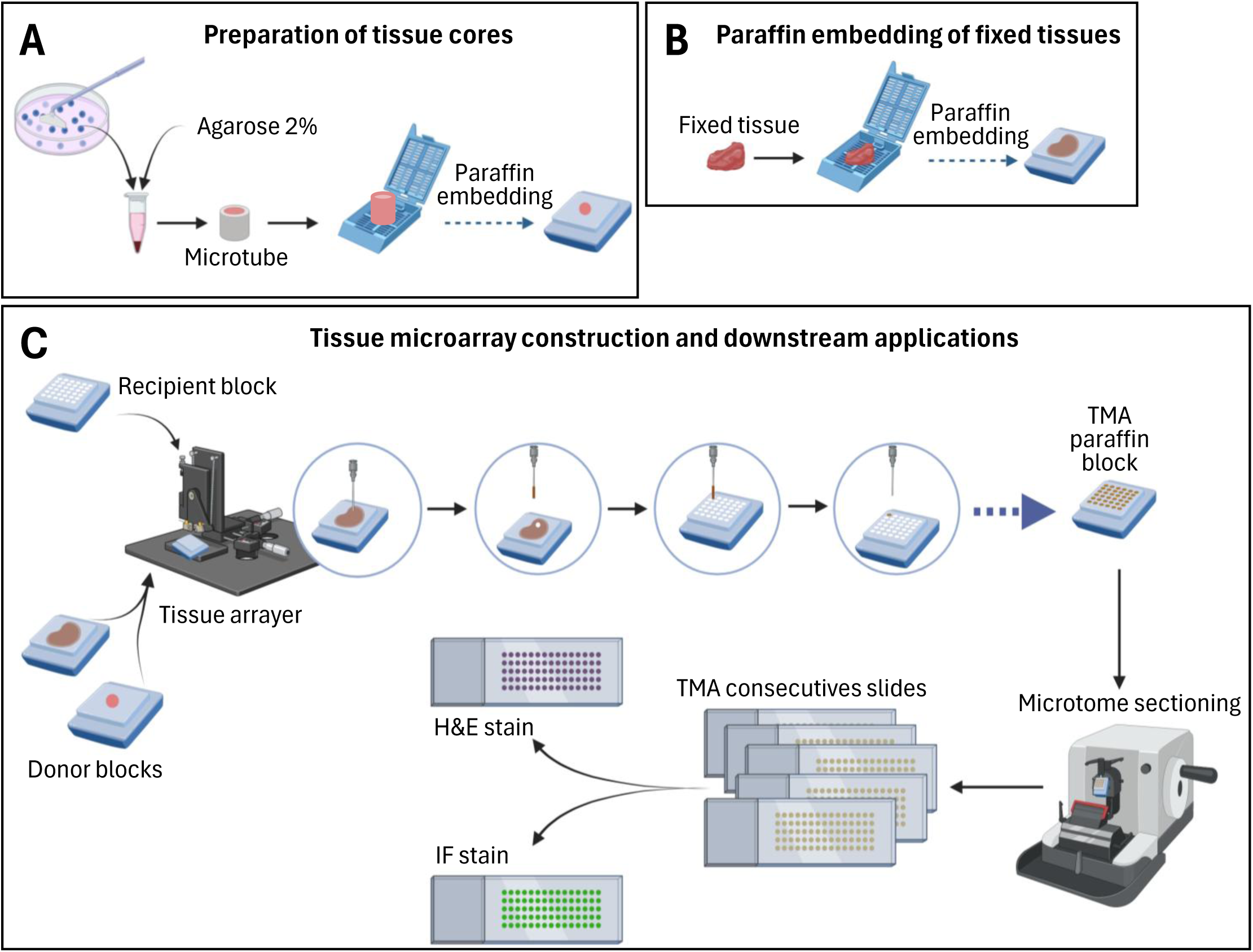
Tissue Microarray preparation. **(A) Preparation of tissue cores.** Tissue samples are first embedded in 2% agarose to stabilize small or irregular specimens prior to processing. Samples are transferred into microtubes and subsequently embedded in paraffin to generate donor blocks. This step ensures preservation of tissue architecture and facilitates precise downstream core extraction. **(B) Paraffin embedding of fixed tissues.** Fixed tissue fragments are directly embedded in paraffin to generate donor blocks suitable for TMA construction. This standard workflow preserves morphological integrity and antigenicity required for histological and multiplex imaging analyses. **(C) Tissue microarray construction and downstream applications.** Cylindrical cores are extracted from donor blocks using a tissue arrayer and precisely inserted into a recipient paraffin block to generate the TMA. This process enables the parallel organization of multiple tissue samples within a single block in a defined spatial layout. The completed TMA block is sectioned by microtomy to produce consecutive slides, ensuring consistent sampling across sections. These sections can then be subjected to hematoxylin and eosin (H&E) staining for morphological assessment or immunofluorescence (IF) staining for multiplex cellular phenotyping. This approach enables high throughput, standardized, and spatially comparable analysis of multiple tissue specimens under identical experimental conditions.

**Supplementary Figure 2.**
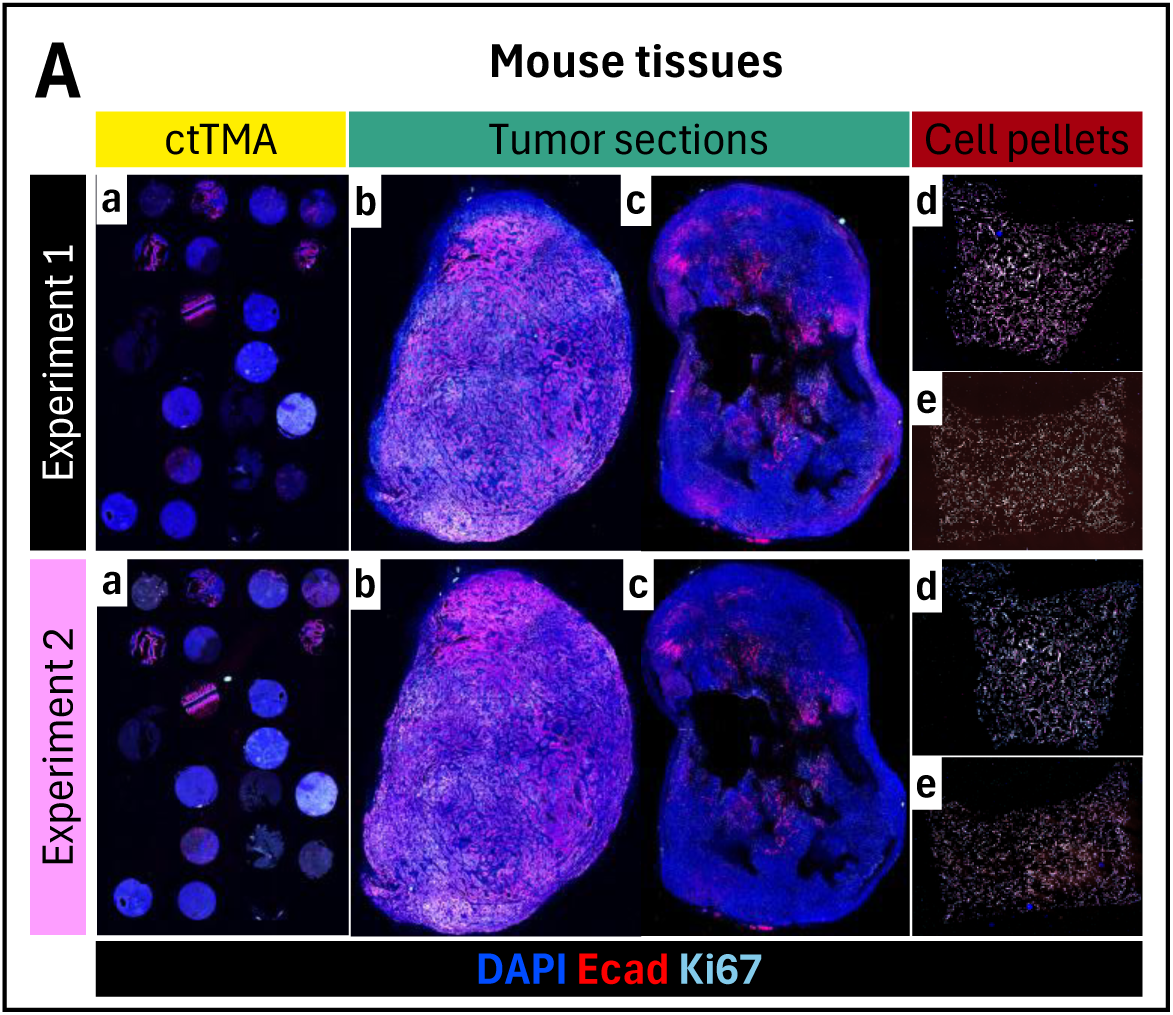
CycIF mouse tissues dual experiments. **(A) CycIF images across independent experiments.** Multiplex immunofluorescence images of mouse tissues processed in two independent experiments (Experiment 1 and Experiment 2), including (a) the reference tissue microarray (ctTMA), (b, c) tumor sections, and (d, e) cell pellets. Representative markers (DAPI, E-cadherin, and Ki67) illustrate nuclear organization, epithelial structures, and proliferative activity, respectively. Across all sample types, comparable staining patterns and tissue architecture are observed between experiments, indicating consistent signal detection and preservation of biological features despite independent processing.

**Supplementary Figure 3.**
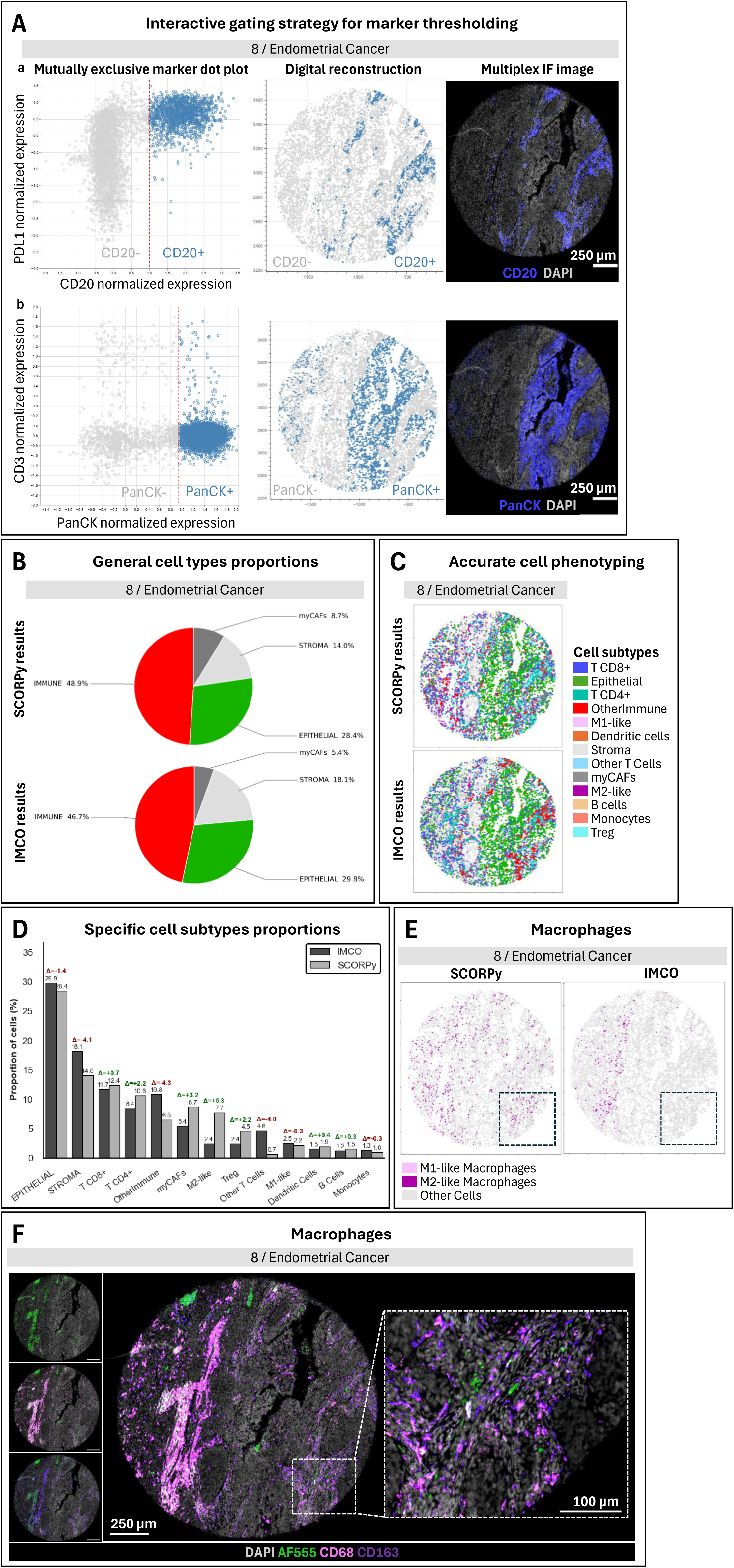
COMET Lunaphore compatibility. **(A) Interactive marker gating and thresholds definition.** Thresholds are interactively adjusted using bivariate scatter plots of normalized marker intensities. In parallel, a synchronized digital tissue reconstruction provides real-time spatial visualization of marker-positive and marker-negative cells, allowing direct validation of gating decisions within their tissue context. (a) CD20 gating is shown using mutually exclusive expression of PDL1. (b) PanCK gating is shown using mutually exclusive expression of CD3. Alongside scatterplots and digital reconstructions, the corresponding multiplex immunofluorescence image is shown for visual reference. Scale bars: 250 µm. **(B) General cell type proportions**. Pie charts showing the distribution of major cell types: epithelial, stromal, and immune cells, obtained in SCORPy software and independently processed and classified in-house IMCO pipeline. **(C) Accurate cell phenotyping**. Spatial distribution of various cell subtypes identified by SCORPy and IMCO pipelines in endometrial cancer TMA core. **(D) Specific cell subtype proportions**. Bar graph comparing the proportions (%) of various cell subtypes as determined by SCORPy (gray bars) and IMCO (black bars) pipelines. Differences between the two methods are indicated with Δ values. **(E) Macrophage populations identification**. Identification of macrophages in endometrial cancer TMA core by SCORPy (left) and IMCO (right) pipelines. The boxed area indicates the region shown at higher magnification in panel **(F)**. **(F) Macrophage populations reference**. Multiplex immunofluorescence images of the same endometrial cancer TMA core showing M1-like macrophages expressing CD68 (light purple), M2- like macrophages expressing CD163 (dark purple), and auto fluorescent cells or artefact (green). The boxed region corresponds to the area highlighted in panel **(E)**. Scale bars, 250 μm for the overview images and 100 μm for the magnified images.

**Supplementary Figure 4.**
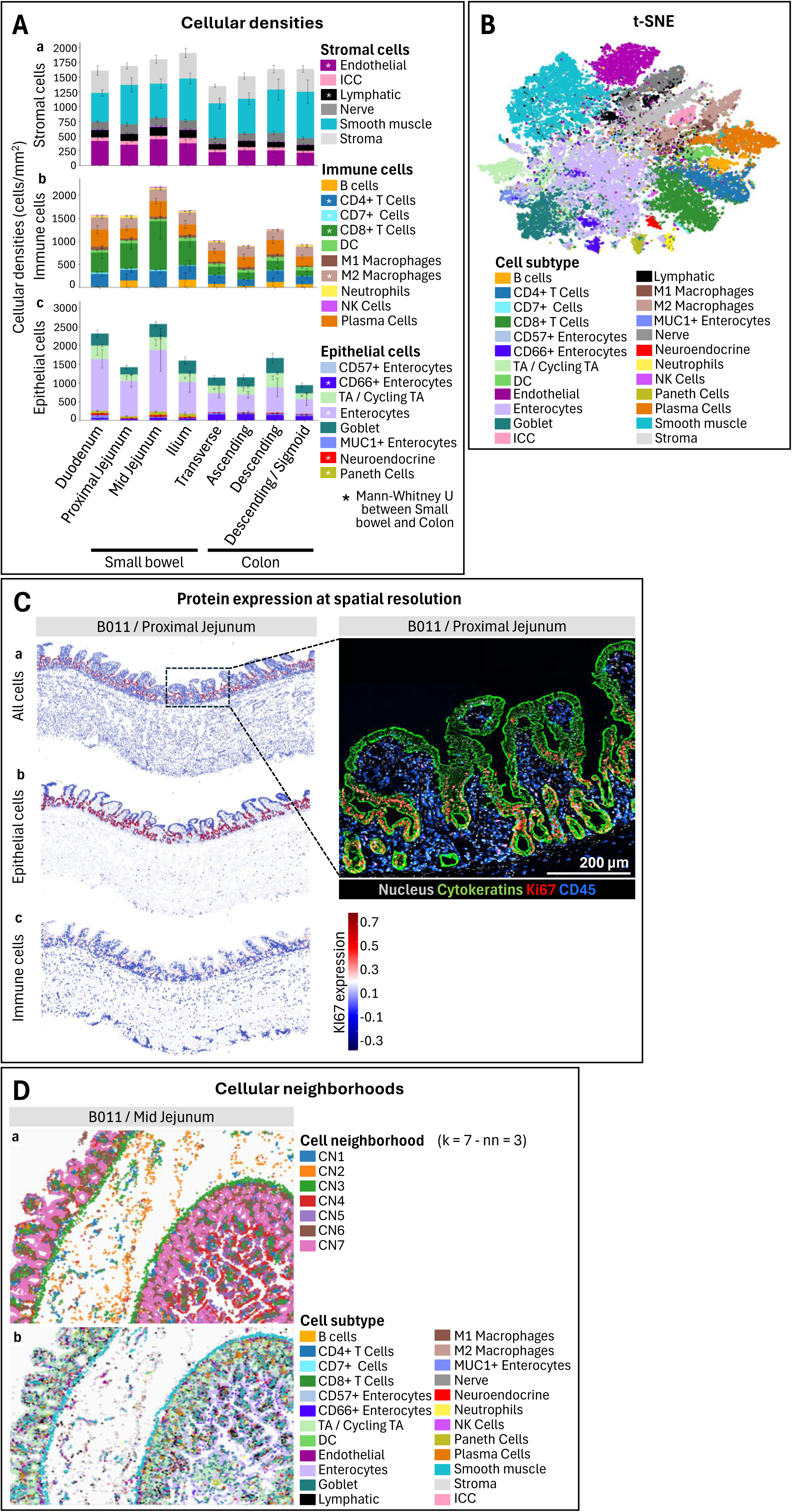
CODEX PhenoCycler compatibility. This figure demonstrates that the analytical framework developed is fully compatible with CODEX PhenoCycler datasets, enabling robust quantification of cell composition, preservation of single-cell phenotypes, and identification of higher-order spatial organization across complex tissues. **(A) Cellular densities across intestinal regions.** Quantification of (a) stromal, (b) immune, and (c) epithelial cell densities across anatomical regions of the intestine, including small bowel and colon segments. Distinct spatial distributions of major cell compartments are observed, reflecting known tissue organization, such as enrichment of epithelial cells in mucosal regions and variation in immune cell abundance along the intestinal tract. Statistical analysis was performed comparing the difference between the small bowel and the colon. **(B) Single-cell phenotypic landscape**. t-SNE projection of all segmented cells colored by cell subtype, revealing clear segregation of major epithelial, immune, and stromal populations. **(C) Spatially resolved protein expression**. Representative images of sample B011 proximal jejunum showing marker expression at single-cell resolution. Spatial maps of (a) all cells, (b) epithelial cells, and (c) immune cells highlight the localization of Ki67-expressing cells within each compartment, enabling visualization of proliferative activity across tissue regions. Multiplex CODEX images with cytokeratins, Ki67 andCD45 further delineate epithelial structures, proliferative zones, and immune cell localization, respectively. The spatial restriction of proliferative (Ki67⁺) cells to specific regions (e.g., crypt-like domains) supports the biological fidelity of the data. Scale bars: 200 µm. **(D) Cellular neighborhood organization.** Spatial clustering of cells into (a) cellular neighborhoods (CN1–CN7) based on local cell-type composition (k = 7, nearest neighbors = 3) in a representative B011 mid-jejunum sample. Distinct neighborhoods correspond to recurring multicellular assemblies, such as epithelial-rich regions, immune-enriched niches, or mixed compartments. Comparison with underlying (b) cell subtype maps shows that these neighborhoods capture higher-order tissue organization beyond individual cell identities, reflecting coordinated spatial patterning. **(E) Visualization of cellular niches.** Unsupervised clustering of spatial neighborhoods defines distinct cellular niches (CN1–CN7). These visualizations highlight the anatomical coherence of niches and their alignment with known intestinal structures, supporting the ability of the analytical framework to extract biologically meaningful spatial domains from CODEX/PhenoCycler data. Clustering was performed with *k* = 7 clusters and *n*ₙₙ = 3 nearest neighbors.

**Supplementary Figure 5.**
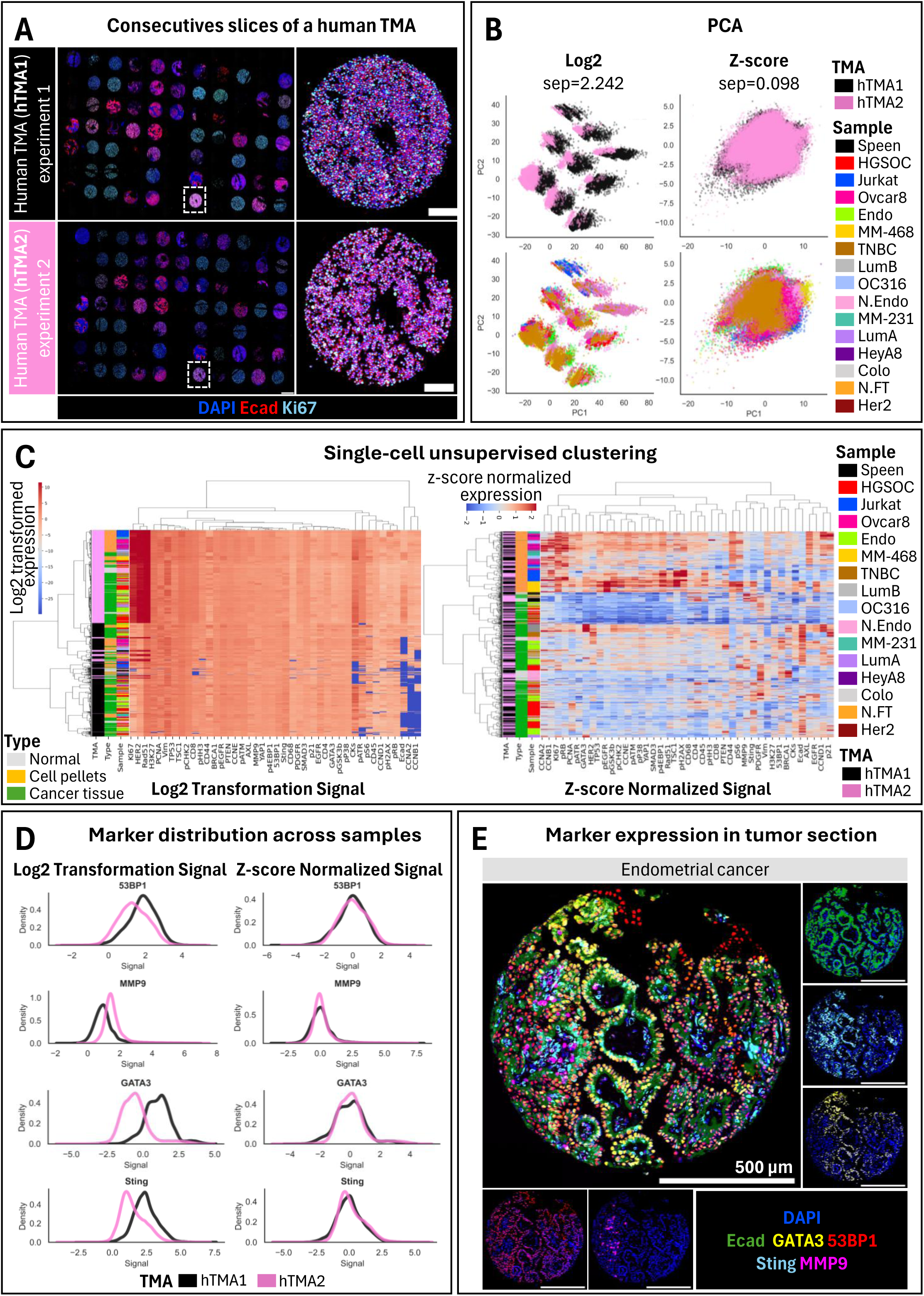
Cross-experiment normalization and batch effect assessment on human tissues. These results extend the findings from Figure 3 to human tissue microarrays, demonstrating that cross-experiment normalization effectively mitigates batch effects while preserving biologically meaningful variation at both the single-cell and tissue levels. **(A) Independent CycIF experiments on human TMAs.** Two consecutive sections of a human tissue microarray (hTMA1 and hTMA2) were processed in two independent experiments. Multiplex immunofluorescence images (DAPI, E-cadherin, Ki67) illustrate consistent tissue architecture and marker localization across experiments, while highlighting potential technical variability in signal intensity prior to normalization. A representative core is shown at higher magnification, corresponding to a cell pellet. **(B) Reduction of batch effects by normalization.** Principal component analysis (PCA) of single-cell data colored by experiment (top) and sample identity (bottom). In log2-transformed data, cells cluster primarily by experiment, indicating strong batch effects (separation = 2.242). Following z-score normalization, separation between experiments is markedly reduced (separation = 0.098), while preserving biologically meaningful grouping by sample type and tumor subtype. **(C) Single-cell clustering preserves biological structure**. Unsupervised hierarchical clustering of single-cell marker expression using log2-transformed and z-score normalized data. While log2- transformed data show clustering influenced by technical variation, Z-score normalization enhances the segregation of biologically relevant groups, including normal tissues, cell pellets, and cancer samples, as well as tumor subtypes, demonstrating improved comparability across experiments. **(D) Marker distribution alignment across experiments**. Density distributions of representative markers (53BP1, MMP9, GATA3, STING) across hTMAs before and after normalization. Log2- transformed signals show experiment-specific shifts, whereas z-score normalization aligns marker distributions between hTMA1 and hTMA2, indicating effective correction of batch effects while maintaining relative expression differences. **(E) Preservation of spatial and biological features after normalization.** Tumor section of endometrial cancer in hTMA1 illustrating marker expression of selected marker (E-cadherin, GATA3, 53BP1, STING, MMP9). Scale bar: 500 µm.,

**Supplementary Figure 6.**
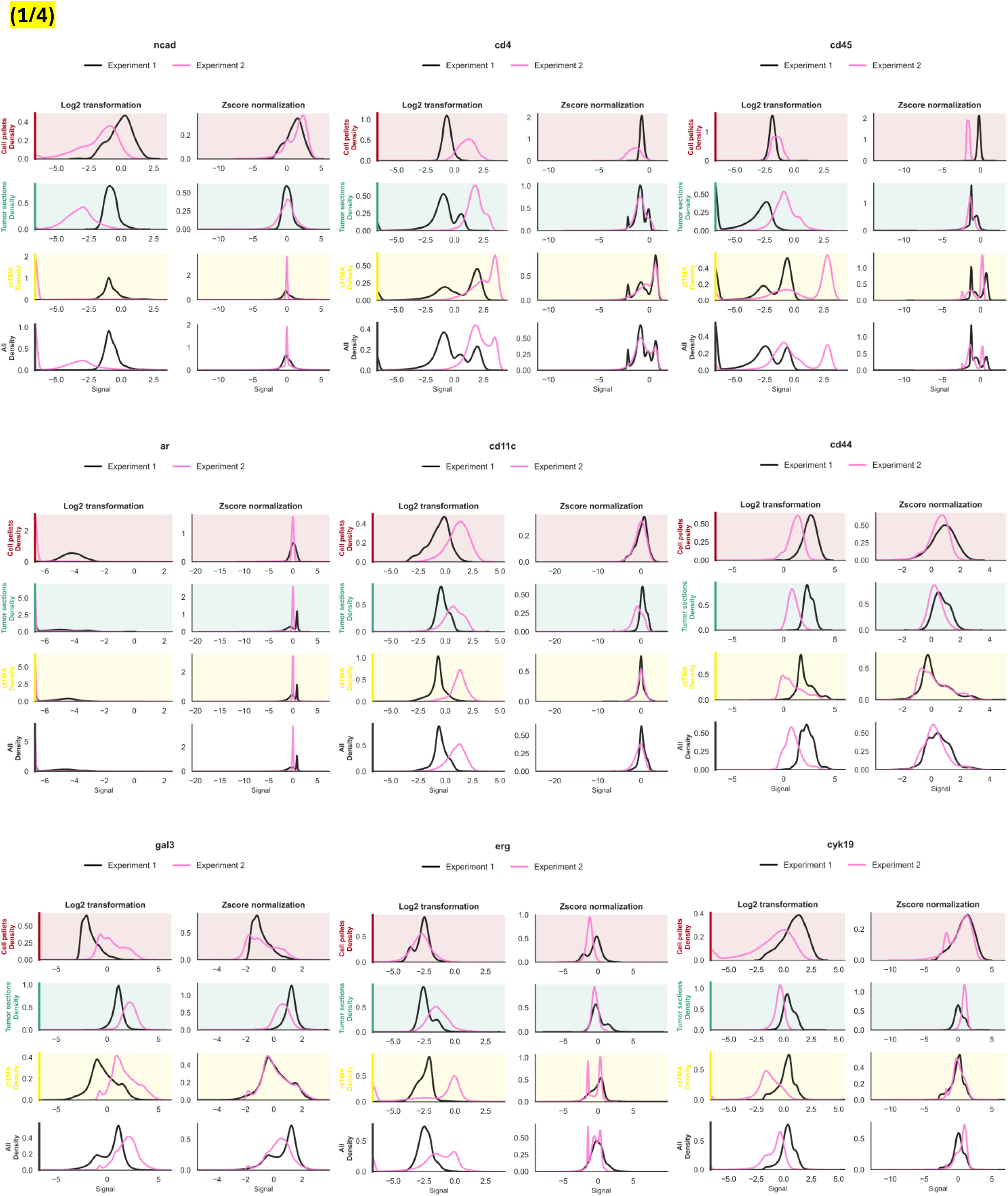
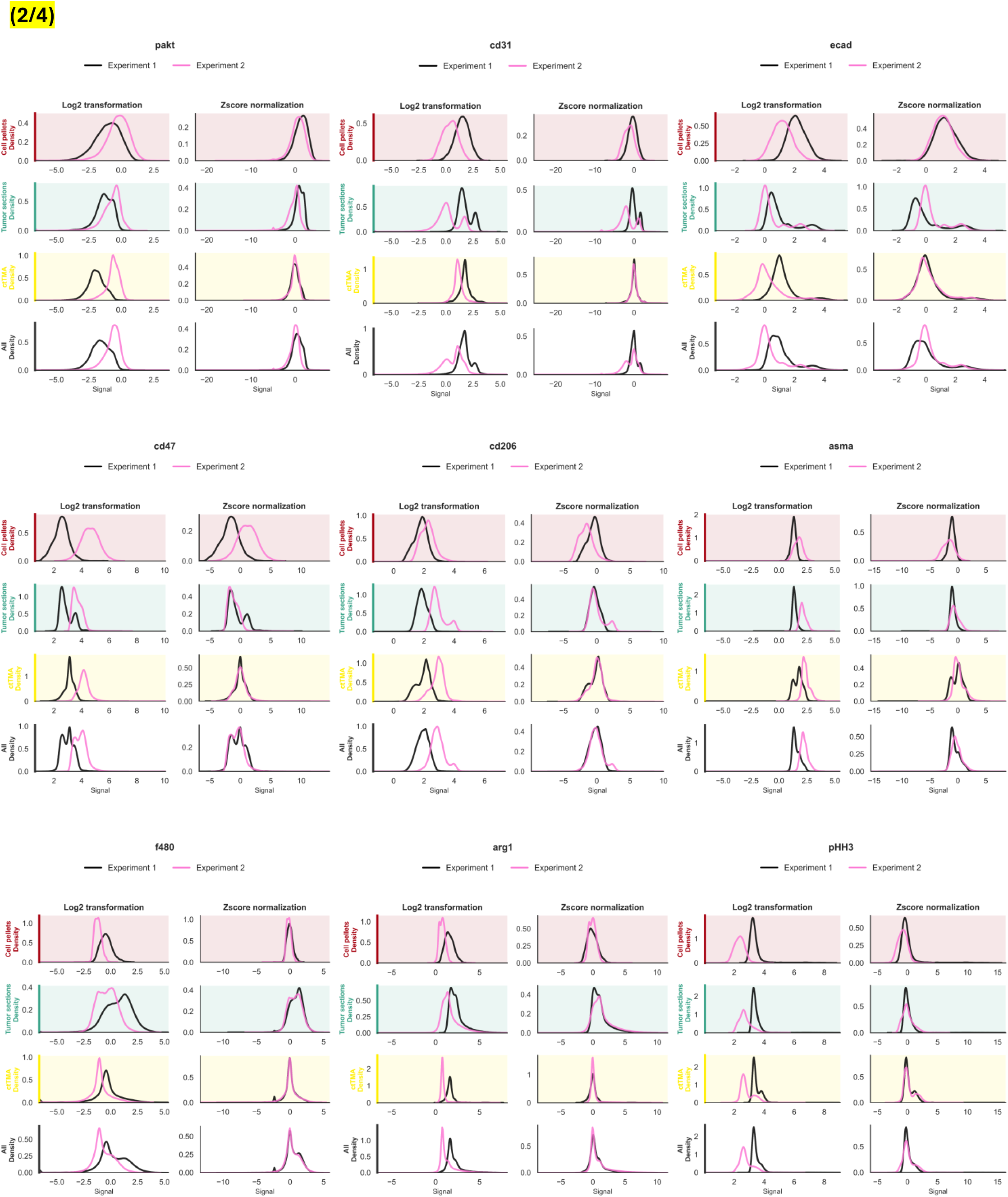
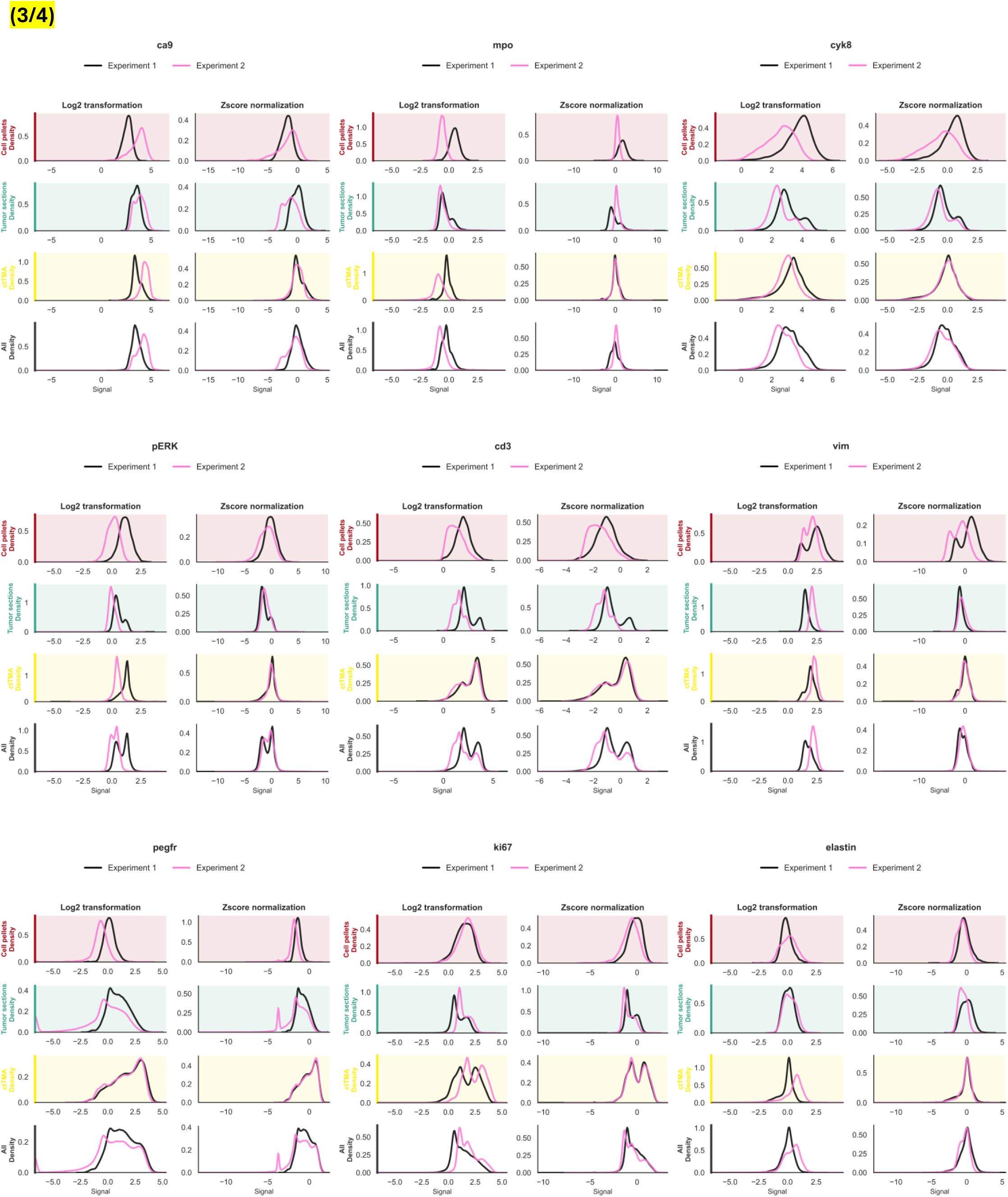
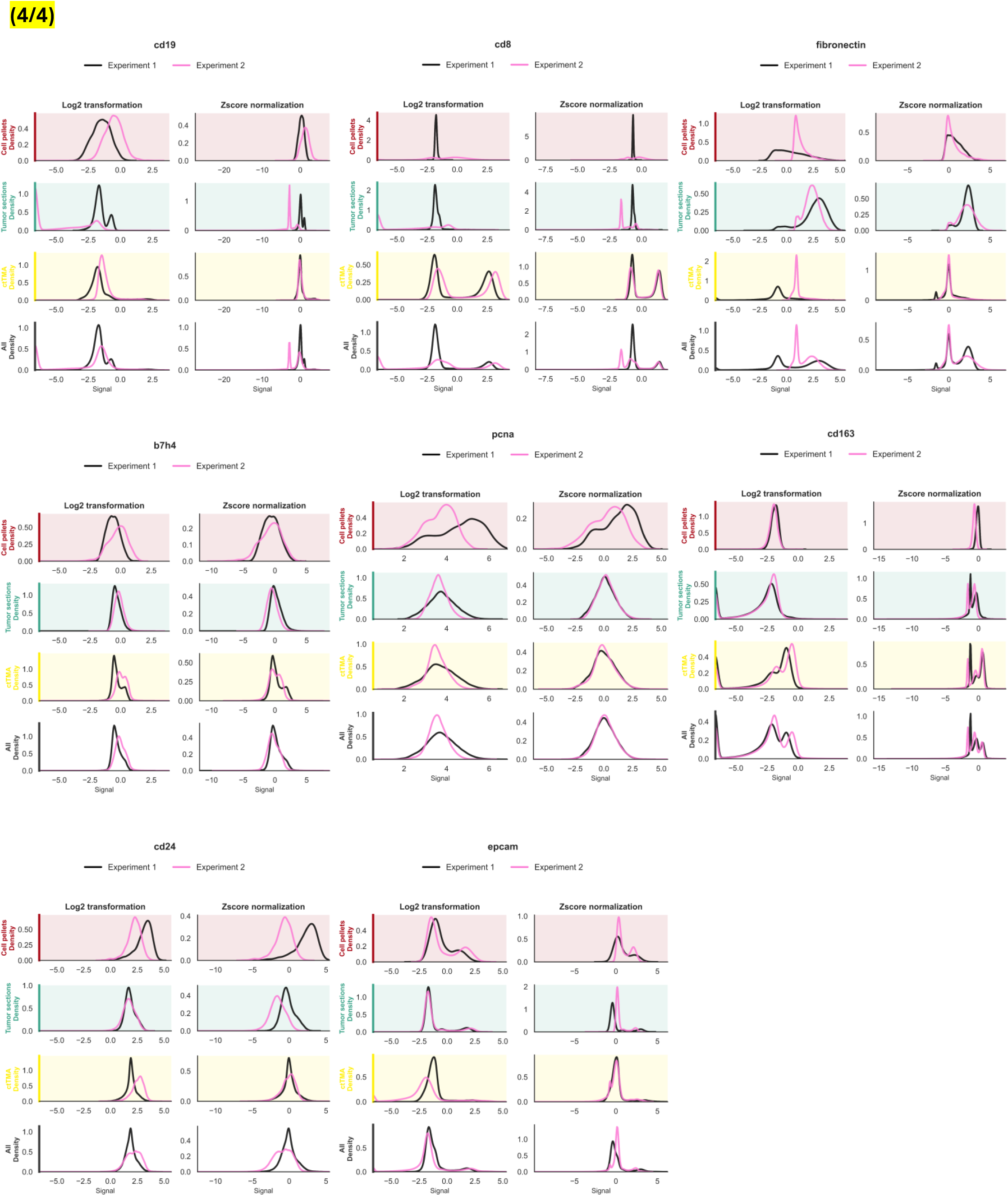
Normalization strategies differentially impact marker intensity distributions across tissues. Density distributions of single-cell marker expression are shown for all proteins in the panel across distinct tissue compartments and samples, comparing two independent CycIF experiments. For each marker, signal distributions are presented after log2 transformation and following z-score normalization. Under log2 transformation, substantial inter-experiment variability is observed, with shifts in signal intensity and distribution shapes depending on the tissue context and marker. In contrast, z-score normalization improves the alignment of distributions between experiments while preserving relative differences across cell types and tissues. These results emphasize the importance of appropriate normalization strategies to enable accurate cross-experiment comparisons while maintaining biologically meaningful variation.

**Supplementary Table 1.**
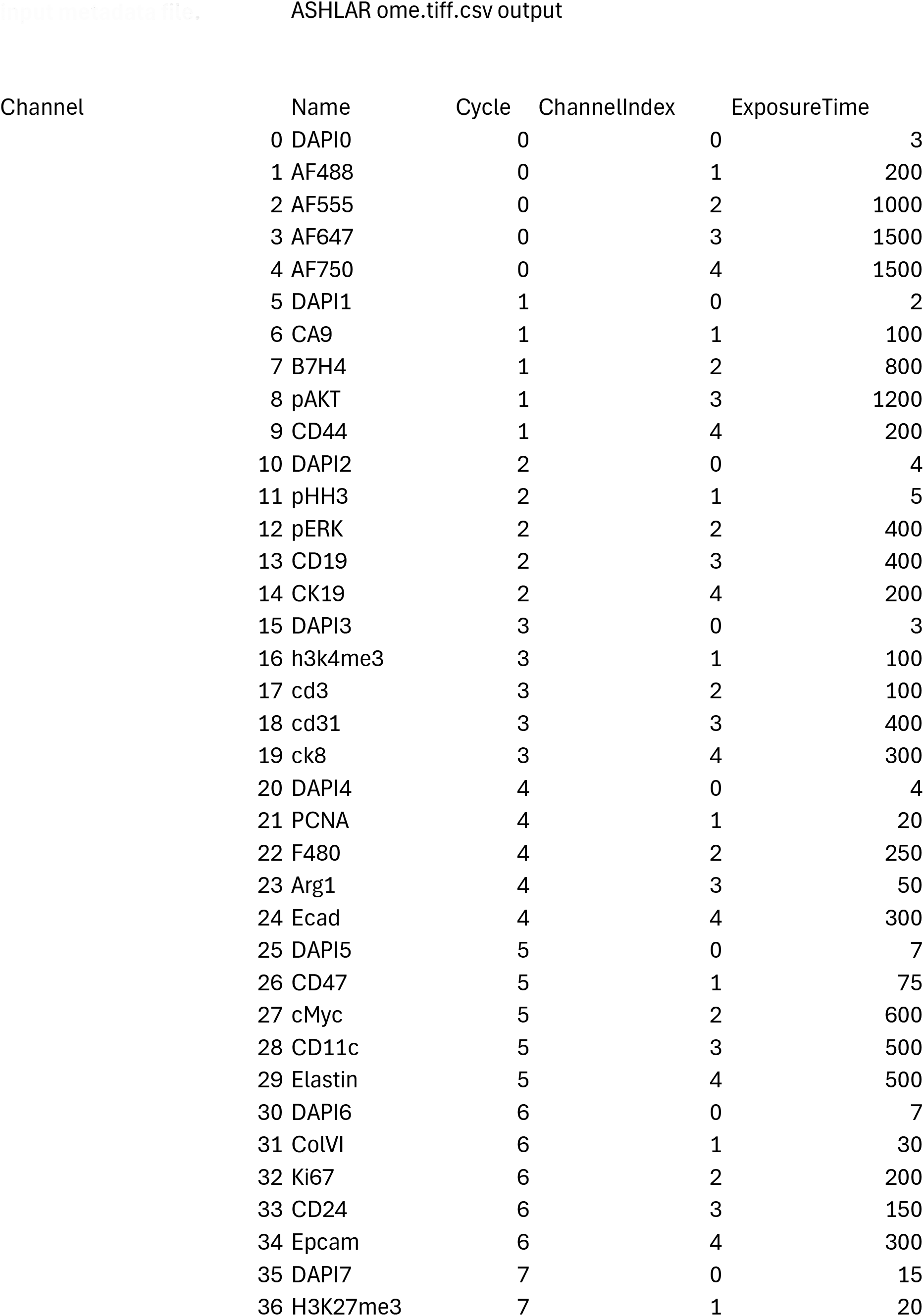
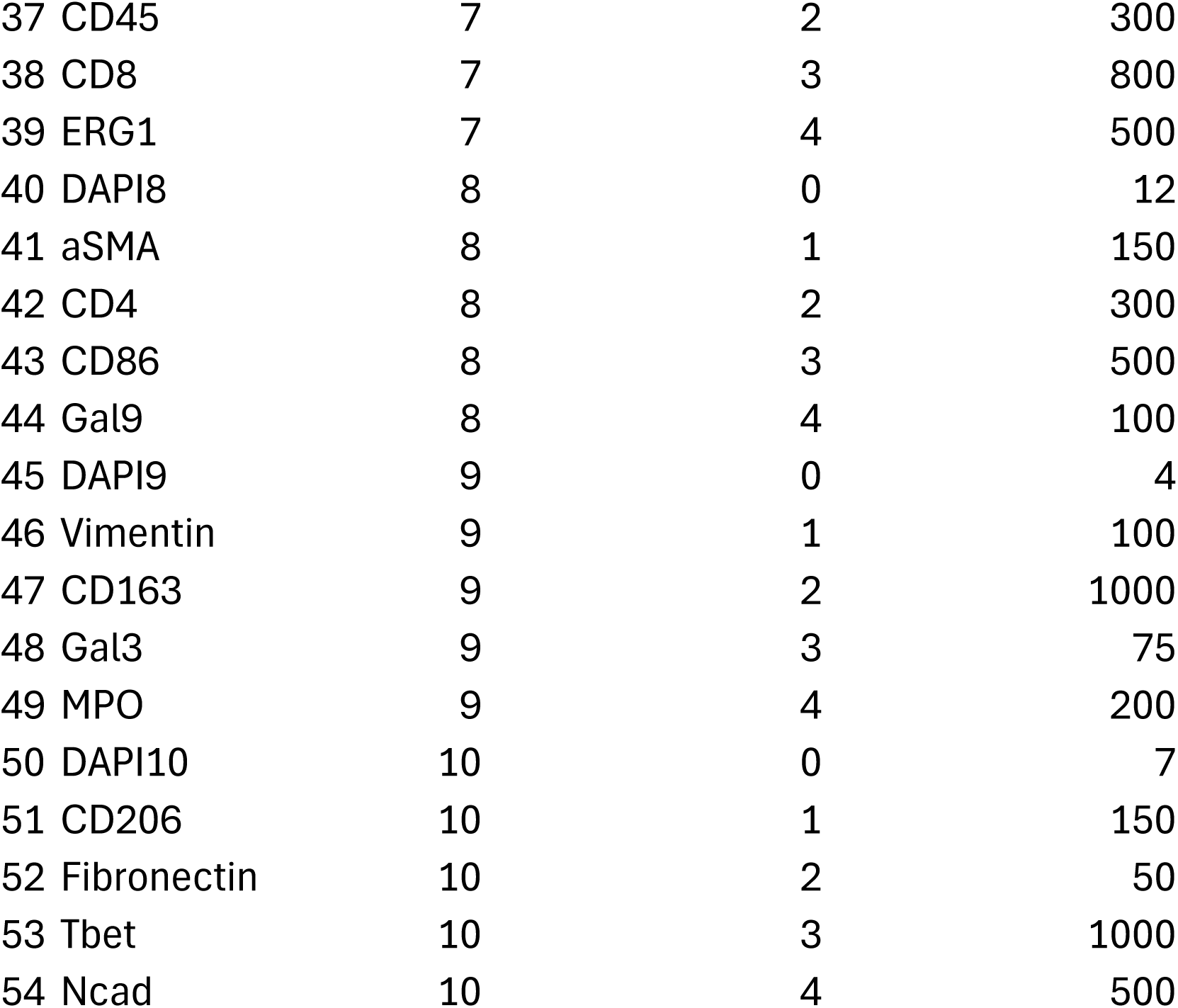
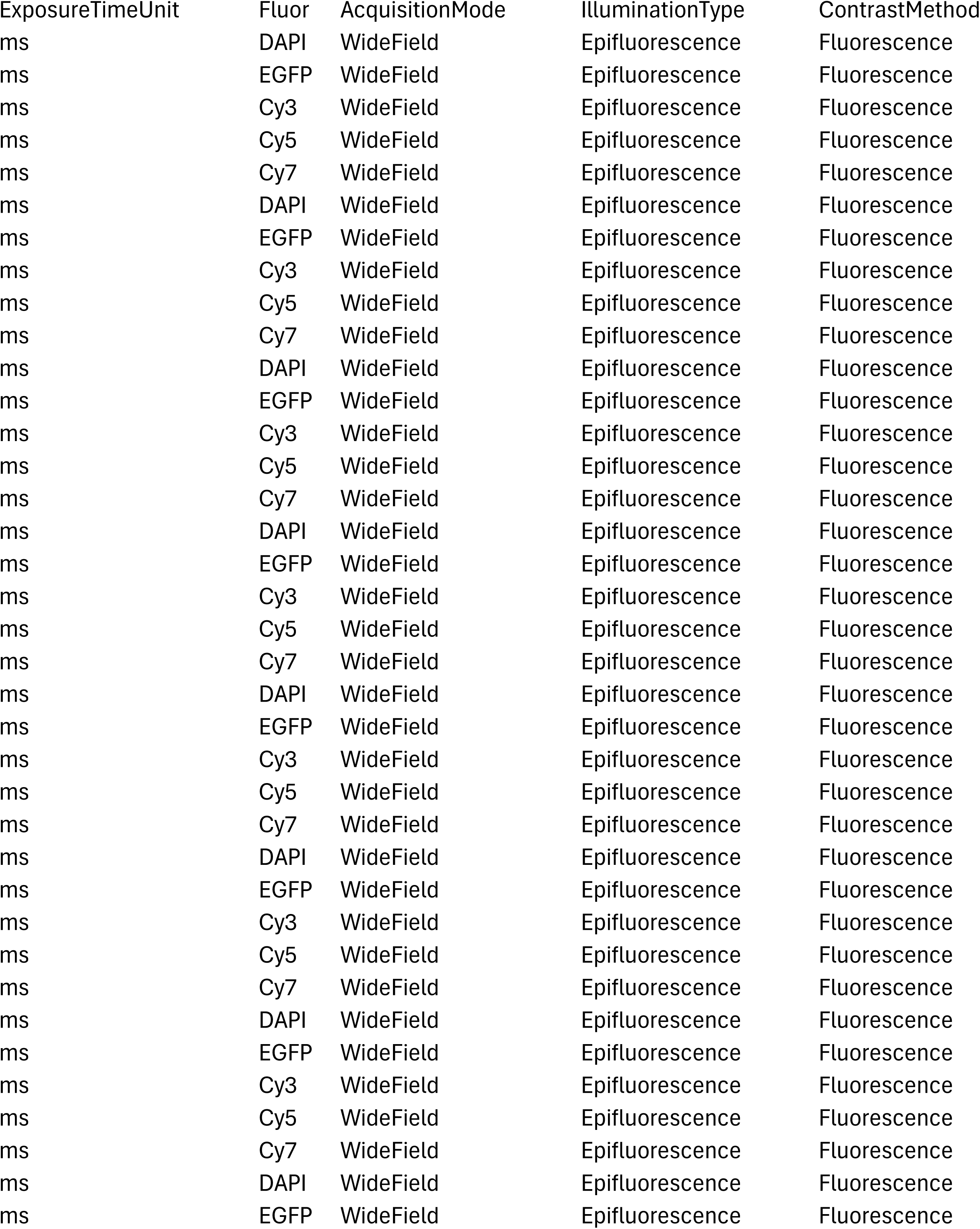
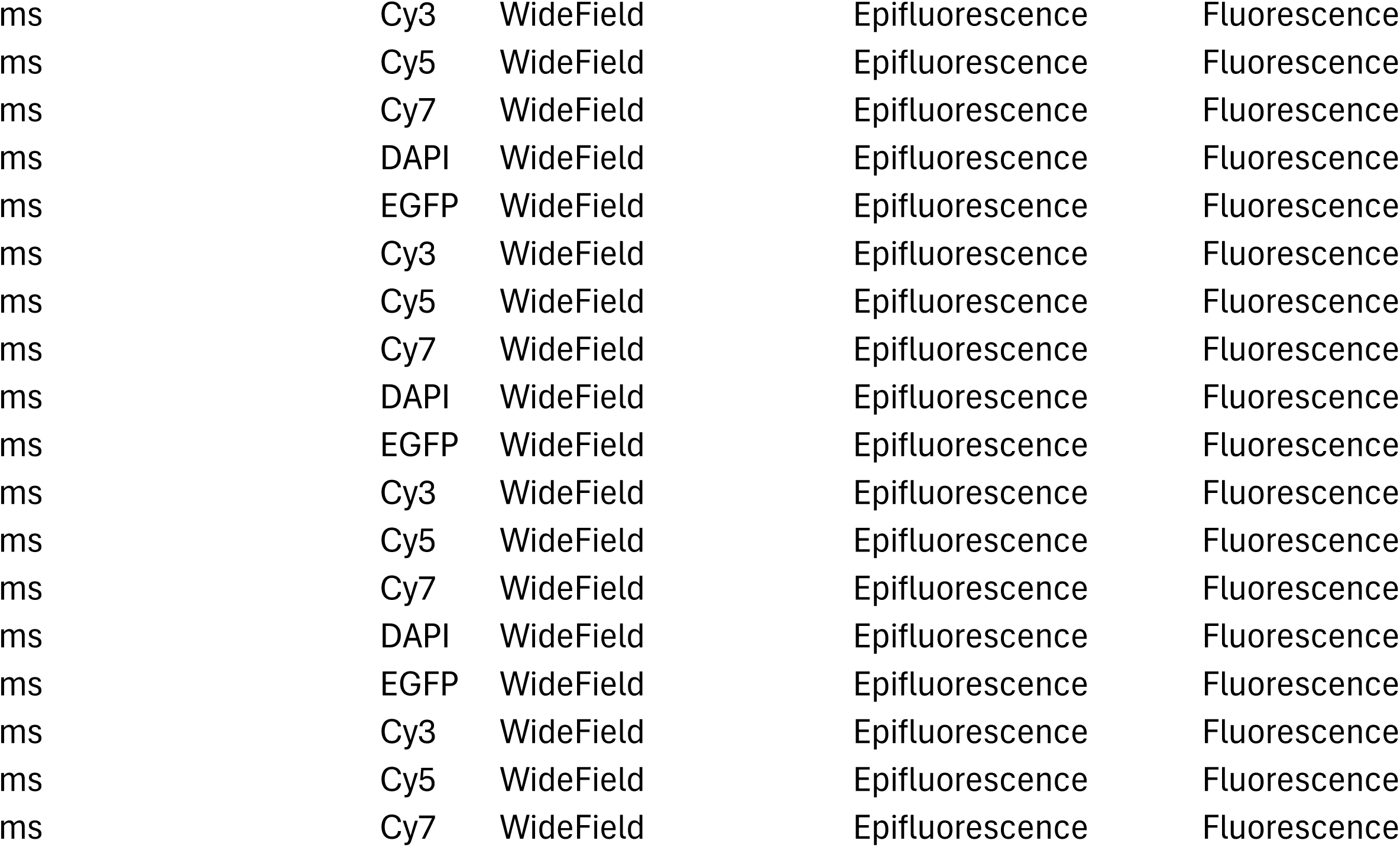
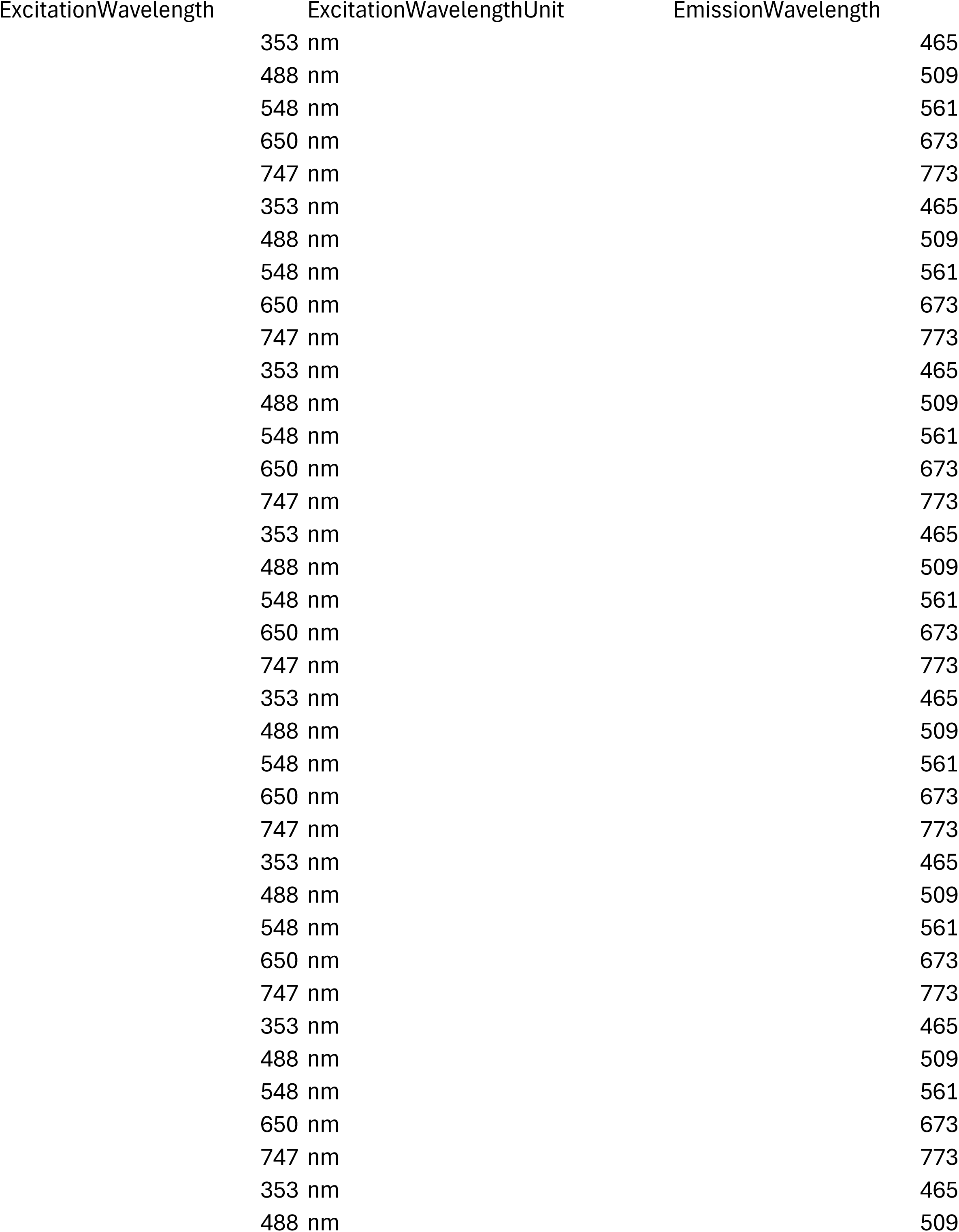
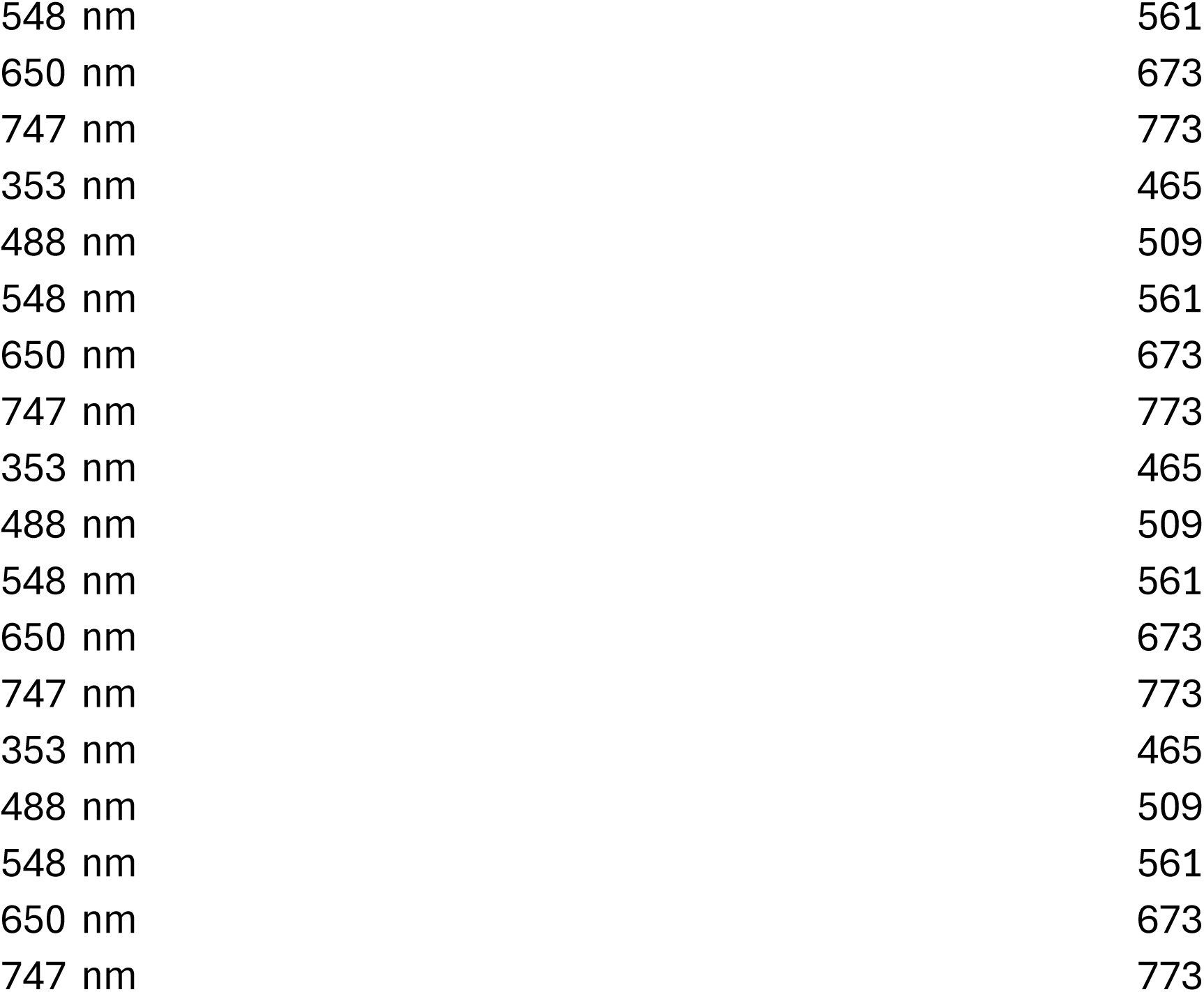
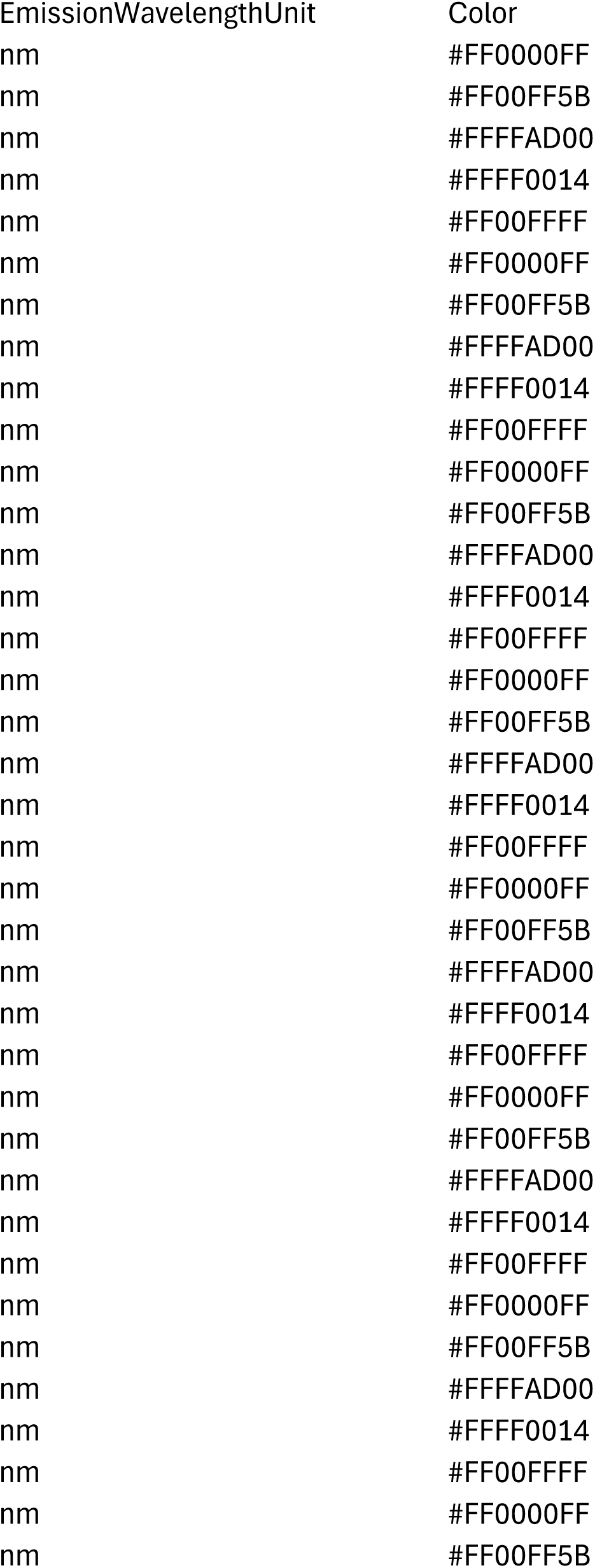
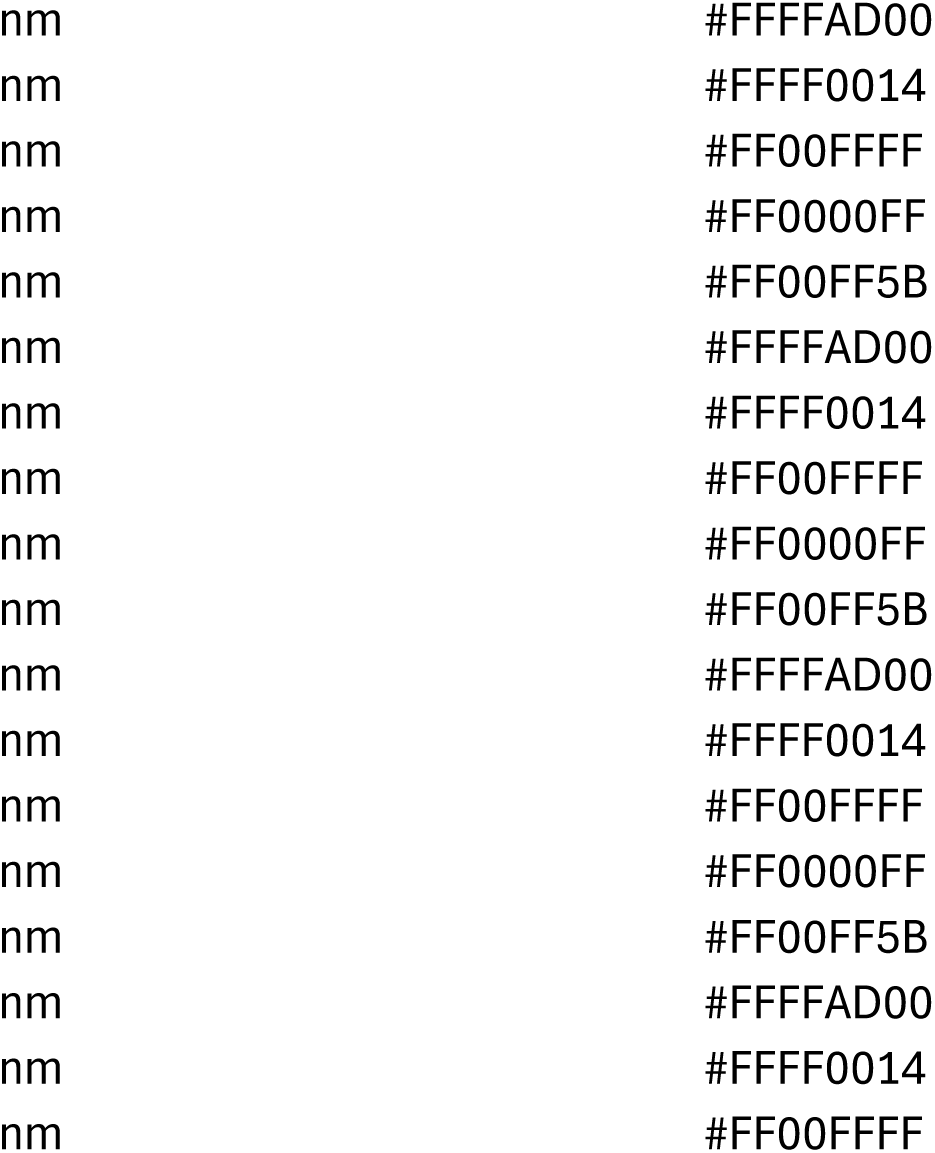
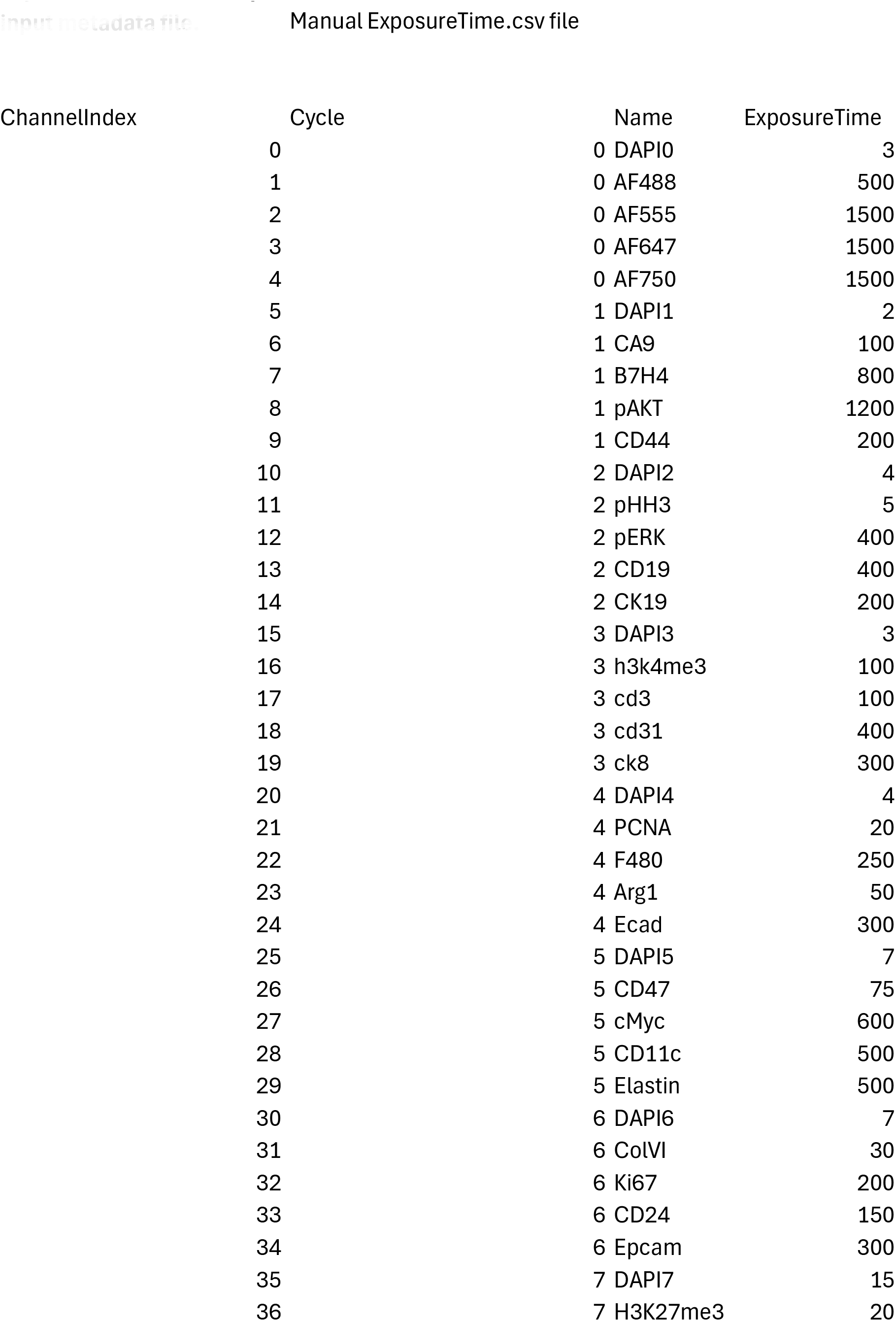
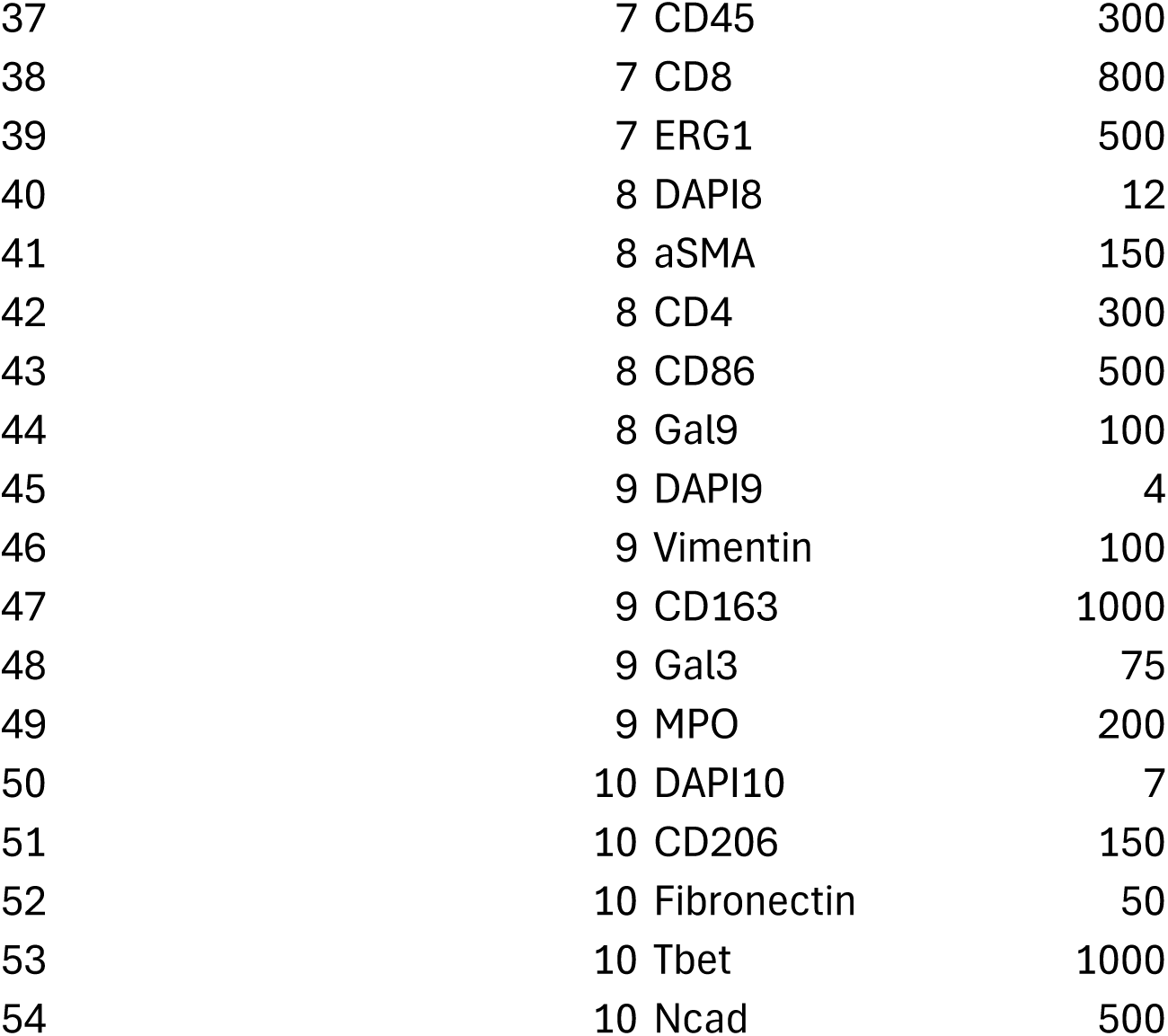
Exposure time example input metadata file.

**Supplementary Table 2.** Column names aliases.

| **Description** | **Standard expected column name** |
| --- | --- |
| X coordinate of the nucleus centroid or cell centroid | **Nuc_X** |
| Y coordinate of the nucleus centroid or cell centroid | **Nuc_Y** |
| Unique identifier for each | **ID** |
| Size/area of the nucleus | **Nucleus_Size** |
| Sample identifier, slide | **Sample_ID** |
| Region of Interest index (for TMA data) | **ROI_index** |
| Classified results : cell | **cell_type** |
| Classified results : cell | **cell_subtype** |

| Aliases |
| --- |
| X, Nuc_X, Nuc X, NucX, Nucleus_X, Nucleus X, x_centroid, X_centroid, centroid_x, Centroid X, Centroid_X, Centroid X um, Centroid X μm, Centroid X μm, Cell X Position, X Position, x |
| Y, Nuc_Y, Nuc Y, NucY, Nucleus_Y, Nucleus Y, Y_centroid, y_centroid, centroid_y, Centroid Y, Centroid_Y, Centroid Y um, Centroid Y μm, Centroid Y μm, Cell Y Position, Y Position, y, Nuc Y Inv, Nucleus Y Inv, Y Inv |
| Cell_ID, Cell Id, CellId, Cell Index, CellIndex, cell_id, ObjectNumber, label, Label, Object ID, Object_ID, |
| Nuc Size, Nuc_Size, NucSize, Nucleus Size, NucleusSize, nucleus_area, Nucleus_Area, Nucleus: Area, |
| Sample ID, SampleID, sample_id, Sample, File, FileName, File Name, file name, Slide, slide_id |
| ROI Index, ROI index, ROIIndex, roi_index, ROI, TMA_core, TMA Core, Core, core_id |
| Cell_Type, CellType, celltype, Cell Type, Classification, classification, Phenotype, phenotype, Cell_Class, |
| Cell_Subtype, CellSubtype, cellsubtype, Cell Subtype, Cell_Subclass, CellSubclass |

| Required | Examples |
| --- | --- |
| Yes | 0, 72.2, -12.333 |
| Yes | 0, 72.2, -12.334 |
| Yes | 0, 1, 2 |
| No | 1, 2, 3 |
| No, but created from filename if not detected | file.csv, slide.csv |
| No, but created and filled with 0 if not detected | 0, 1, 2 |
| Yes, if import from cassified module, no otherwise | Immune, stroma |
| No | T CD8+, B Cells, Stroma |

**Supplementary Table 3.** Antibodies information.

ctTMA CyclIF Panel
| R0 | AF488 | AF555 | AF647 | AF750 |
| --- | --- | --- | --- | --- |
| R1 | CA9 | B7H4 | pAKT | CD44 |
| R2 | pHH3 | pERK | CD19 | CK19 |
| R3 | CD47 | CD3 | CD31 | CK8 |
| R4 | PCNA | f4/80 | Arg1 | Ecad |
| R5 | H3K4me3 | cMyc | CD11c | Elastin |
| R6 | ColVI | Ki67 | CD24 | EpCam |
| R7 | H3k27me3 | CD45 | CD86 | ERG1 |
| R8 | aSMA | CD4 | CD8 | Gal9 |
| R9 | Vimentin | CD163 | Gal3 | MPO |
| R10 | CD206 | Fibronectin | Tbet | Ncad |

**Supplementary Table 3.****Antibodies information.** Antibodies informations
| Target | Clone | Company | Catalog number |
| --- | --- | --- | --- |
| Arg1 | EPR22033-369 | Abcam | ab307468 |
| aSMA | EPR5368 | Abcam | ab202295 |
| B7H4 | EPR23665-20 | Abcam | ab252438 |
| CA9 | EPR23055-5 | Abcam | ab309504 |
| CD11c | D1V9Y | CST | 97585S |
| CD163 | EPR19518 | Abcam | ab281746 |
| CD19 | EPR23174-145 | Abcam | ab314289 |
| CD206 | 2A6A10 | Proteintech Group Inc. | CL488-60143 |
| CD24 | poly | NOVUS | NBP1-46390 |
| CD3 | D7A6E | CST | 57869S |
| CD31 | EPR17259 | Abcam | ab305210 |
| CD4 | EPR19514 | Abcam | ab313727 |
| CD44 | poly | Abcam | ab157107 |
| CD45 | 6O19 | Protein Tech | CL555-80297 |
| CD47 | SP279 | Abcam | ab290989 |
| CD8 | EPR21769 | Abcam | ab237365 |
| CD86 | C86/1146 | Abcam | ab220188 |
| cMyc | Y69 | Abcam | ab201780 |
| ColVI | EPR17072 | Abcam | ab200429 |
| CK19 | A53-B/A2 | Biolegend | 628502 |
| CK8 | 1E8 | Biolegend | 904804 |
| Ecad | poly | Sino biological | 50671-RP01 |
| Elastin | poly | Signalway antibody | C45078-AF750 |
| EpCam | EPR20533-63 | Abcam | ab221552 |
| ERG1 | EPR3864 | Abcam | ab92513 |
| f4/80 | D2S9R | CST | 99651 |
| Fibronectin | F14 | Abcam | ab45688 |
| Gal3 | M3/38 | Biolegend | 125408 |
| Gal9 | poly | Abcam | ab69630 |
| H3K27me3 | C36B11 | CST | 5499S |
| H3K4me3 | poly | MilliporeSigma | 07-473-AF488 |
| Ki67 | 8D5 | CST | 9449S |
| MPO | EPR17996 | Abcam | ab188211 |
| Ncad | poly | Proteintech | 22018-1-AP |
| pAKT | D9E | CST | 4075S |
| PCNA | PC10 | CST | 8580S |
| pERK | poly | R&D | AF1018 |
| pHH3 | D2C8 | CST | 3465S |
| Tbet | EPR9302 | Abcam | ab225206 |
| Vimentin | D21H3 | CST | 9854S |

